# Targeting FGFR signaling overcomes therapeutic resistance and immune evasion in oncogenic PIK3CA-driven serous-like endometrial cancer

**DOI:** 10.64898/2026.02.16.706009

**Authors:** Xin Cheng, Changli Qian, Erica Holdridge, Guruprasad Ananda, Tao Jiang, Yadong Zhang, Jing Ni, Shaozhen Xie, Hao Gu, Renlei Ji, Elena V Ivanova, Marisa R Nucci, Zhe Wang, Kaifu Chen, Bose Kochupurakkal, Gordon J. Freeman, Geoffrey I. Shapiro, Joyce Liu, Panagiotis A. Konstantinopoulos, Ursula Matulonis, Jean J. Zhao

## Abstract

Serous endometrial cancer (SEC) is an aggressive subtype of endometrial cancer (EC) with poor prognosis and limited treatment options. Here, we developed a clinically relevant, immunocompetent serous-like mouse model incorporating oncogenic *PIK3CA* mutation, *Trp53* loss, and *MYC* overexpression. Using this model together with human EC cell lines, patient-derived organoids (PDOs), xenografts, and patient datasets, we investigated mechanisms underlying resistance to PI3Kα-targeted therapy. Single-cell profiling reveals that FGFR1/2 upregulation associates with intrinsic resistance, whereas FGFR3 characterizes acquired resistance. Dual FGFR and PI3Kα inhibition produced superior tumor control compared with either agent alone. Mechanistically, FGFR signaling promotes immune evasion by downregulating MHC-I/HLA-mediated antigen presentation and enriching M2-type tumor-associated macrophages. FGFR inhibition reversed these changes and synergized with anti-PD-1 therapy to enhance antitumor immune responses and establish durable immune memory. Collectively, these findings identify FGFR signaling as a key driver of therapeutic resistance and immune escape in SEC and support FGFR-targeted combination strategies.

## INTRODUCTION

Endometrial cancer (EC) is the most common gynecologic malignancy in the United States, with incidence and mortality rates continuing to rise ^1^. SEC, a particularly aggressive subtype, accounts for nearly 40% of EC-related deaths ^2^. SEC is characterized by frequent TP53 mutations (∼90%) and a “serous-like” molecular profile with extensive copy-number alterations, and is associated with markedly reduced overall survival ^3^. SEC also exhibits frequent alterations in the PI3K pathway, including mutations in PIK3CA (48%) and PIK3R1 (13.3%), as well as additional aberrations such as FGFR overexpression or amplification and MYC amplification (44%) ^4, 5^.

Current treatment options for SEC remain limited. Standard chemotherapy yields response rates of only 10-15% and a median overall survival of 8-12 months in advanced disease ^6, 7^. Immune checkpoint inhibitor (ICI) monotherapy shows similarly poor response rates (2.8-6.3%) ^8, 9, 10^. Lenvatinib, a multiple tyrosine kinase inhibitor targeting FGFR1-4, VEGFR1-3, RET, and KIT, has demonstrated clinical activity in phase II/III trials ^11, 12^. Although the FDA-approved pembrolizumab-lenvatinib combination improves response rates to ∼38%, treatment is often limited by significant toxicity, and approximately 40% of patients eventually experience disease progression. The absence of predictive biomarkers further complicates treatment selection ^13, 14, 15^.

The phosphoinositide-3 kinase (PI3K) pathway is a crucial therapeutic target in SEC, with mutations present in 93% of EC cases ^4, 16^. Among these, PIK3CA mutations, predominantly in the helical region (E545K, E542K) and the kinase domain (H1047R), lead to the constitutive activation of PI3Kα, promoting tumorigenesis and disease progression ^17, 18, 19^. Although several PI3Kα-selective inhibitors, including alpelisib (BYL719), copanlisib, inavolisib (GDC-0077), and STX-478, have shown clinical efficacy in advanced solid tumors and lymphoma ^20, 21, 22, 23^, PI3K inhibitors in endometrial cancer including serous endometrial cancers have demonstrated limited activity ^24, 25, 26^. Several mechanisms of resistance to PIK3CA-targeted therapies have been reported, including feedback activation of AXL/EGFR/PKC signaling, compensatory activation of PI3Kβ in PTEN-deficient tumors, and reactivation of parallel MAPK and mTORC1 pathways ^27, 28, 29, 30^. Additional studies revealed metabolic and kinase rewiring involving PIM1 upregulation and insulin/IGF feedback loops, that restore AKT-mTOR signaling and promote tumor survival despite PI3K inhibition ^31, 32^. However, the specific mechanisms of resistance in EC remain poorly understood.

In this study, we developed a clinically relevant immunocompetent transgenic mouse model of serous-like endometrial cancer driven by PIK3CA mutation, TP53 loss, and MYC overexpression. This model enables systematic investigation of PI3Kα-targeted therapy resistance within an immunocompetent SEC context. Using this system, we identified FGFR signaling as a key driver of resistance to PI3Kα-targeted therapy and demonstrate that combined FGFR and PI3Kα inhibition, particularly when integrated with immune checkpoint blockade, elicits enhanced antitumor efficacy. Together, these findings establish a mechanistic and therapeutic framework for targeting FGFR-mediated resistance and immune evasion in SEC.

## Results

### Generation of a syngeneic orthotopic serous-like endometrial cancer model driven by the PIK3CA-H1047R mutation

To investigate the role of PI3K pathway activation in SEC development and resistance to PI3Kα inhibition, we generated a syngeneic orthotopic genetically engineered mouse model (GEMM) of SEC that combines oncogenic PIK3CA mutation, TP53 loss and MYC overexpression (termed APM), incorporating common genetic alterations observed in human SEC^3, 4, 5^. This APM model was initially established by isolating uterine epithelial cells from FVB mice carrying three genetic elements: Rosa-lsl-rtTA, floxed-Trp53, and a tetracycline (tet)-inducible PIK3CA^H1047R^-IRES-luciferase transgene ^33^. These cells were then modified with adenoviral Cre and lentiviral mouse Myc before being orthotopically transplanted into the uterine wall of FVB female recipients (**Fig. 1a**). Following doxycycline administration, transplanted mice developed *de novo* primary APM uterine tumors within 3-5 months with full penetrance. In contrast, mice not treated with doxycycline remained tumor-free, demonstrating that PIK3CA-H1047R expression is required for tumor development (**Fig. 1b**). RT-qPCR and western blot analysis confirmed that doxycycline treatment significantly induced PIK3CA^H1047R^ expression and activated downstream PI3K signaling. While endogenous murine Pik3ca expression remained unaffected (**Fig. 1c and Supplementary Fig. 1a, b**). The tet-inducible PIK3CA-H1047R transgene was coupled to a luciferase reporter for in vivo monitoring of transgene expression (**Supplementary Fig. 1c**).

**Figure 1.**
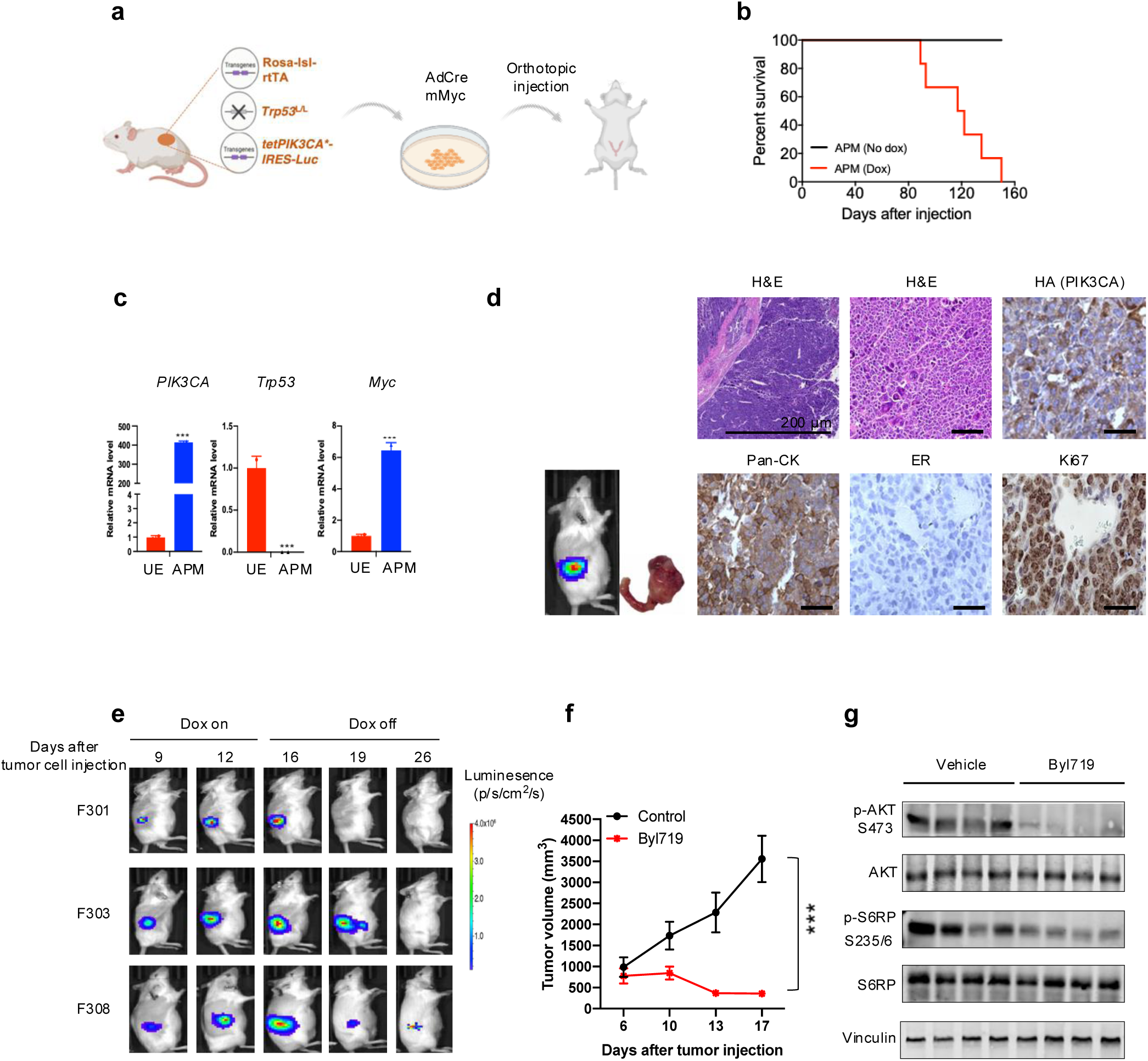
Characterization of the PIK3CA^H1047R^; Trp53^-/-^; Myc (APM) mouse model. **(a)** Schematic illustrating the strategy used to generate the APM transgenic model. **(b)** Kaplan-Meier survival curves comparing APM mice with doxycycline-naïve transgenic controls (n=6 per group). **(c)** RT-qPCR analysis of PIK3CA^H1047R^ transgene, Trp53, and Myc expression (n=3 biologically independent samples). **(d)** Representative histology of primary APM tumors showing serous-like morphology (scale bar, 50 μm). **(e)** Representative in vivo bioluminescence imaging showing regression of primary tumors following doxycycline withdrawal. **(f)** Tumor volume changes in APM allografts treated with BYL719 (20 mg/kg) or vehicle (n=8 each group). **(g)** Immunoblot analysis of tumor lysates from mice treated with BYL719 or vehicle.

These primary tumors exhibited histological characteristics of SEC, including endometrial atrophy, papillary/micropapillary architecture with slit-like spaces, marked nuclear atypia, necrosis, and loss of TP53 expression. Tumor cells overexpressed PIK3CA and MYC, were estrogen receptor (ER)-negative, and co-expressed epithelial markers (**Fig. 1d**). We next benchmarked the APM model against a well-established PTEN-loss-driven EC GEMM, which has been extensively characterized across multiple studies and classified as a Type I-like, endometrioid EC ^34, 35, 36^. RNA-seq-based gene set enrichment analysis (GSEA) revealed that APM tumors exhibited strong enrichment of G2M checkpoint activation, DNA replication stress-associated programs, E2F targets, and MYC target pathways relative to the PTEN-loss model ^36^, consistent with molecular features reported for human SEC ^4^ (**Supplementary Fig. 1d**). Copy-number variation (CNV) analysis further demonstrated extensive genomic instability in APM tumors, with a fraction of genome altered (FGA) of 0.33, comparable to human SEC tumors (median FGA = 0.41) and markedly higher than that observed in non-serous TCGA endometrial cancer (median FGA =0.07) (**Supplementary Fig. 1e**).

To create a more time-efficient and experimentally tractable system for mechanistic and therapeutic studies, we then derived primary tumor cells from these *de novo* APM tumors and orthotopically injected them into the uterine wall of FVB female recipients. Upon doxycycline administration, tumors developed in all mice within one month, providing a reliable experimental model with predictable growth kinetics (**Supplementary Fig. 1f**). We assessed tumor dependency by withdrawing doxycycline from tumor-bearing mice, which resulted in rapid reduction of PIK3CA^H1047R^ expression and downstream signaling pathway activation (**Fig. 1e and Supplementary Fig. 1g**). Tumor cells underwent apoptosis within 24-48 hours, followed by tumor regression within one week (**Supplementary Fig. 1h**), indicating their dependence on PIK3CA-H1047R.

Finally, to assess pharmacologic inhibition, syngeneic transplants were treated with the PI3Kα-selective inhibitors BYL719 (alpelisib) and GDC-0077 (inavolisib). Consistent with genetic inactivation, both agents markedly suppressed tumor growth (**Fig. 1f and Supplementary Fig. 1i, j**) and reduced phosphorylation of AKT and S6RP (**Fig. 1g**), confirming that APM tumor growth is dependent on sustained PIK3CA-H1047R signaling.

### Tumors recurring after PIK3CA withdrawal exhibit FGFR-dependent proliferation

To model tumor recurrence, which commonly arises from therapeutic resistance in a small population of tumor cells and can manifest as both local and distant relapse ^37^, we withdrew doxycycline from a cohort of 14 tumor-bearing mice and monitored them for several months. About one-third (5/14) of tumors underwent complete regression within one month and did not recur (**Fig. 2a**), indicating that these tumors remained dependent on PIK3CA^H1047R^ signaling for their survival. In contrast, approximately two-thirds of tumors (9/14) initially regressed but later resumed growth despite the continued absence of doxycycline (**Fig. 2a and Supplementary Fig. 2a**). Since control mice without doxycycline induction never developed tumors, these recurrent tumors must have originated from resistant clones within the original tumors.

**Figure 2.**
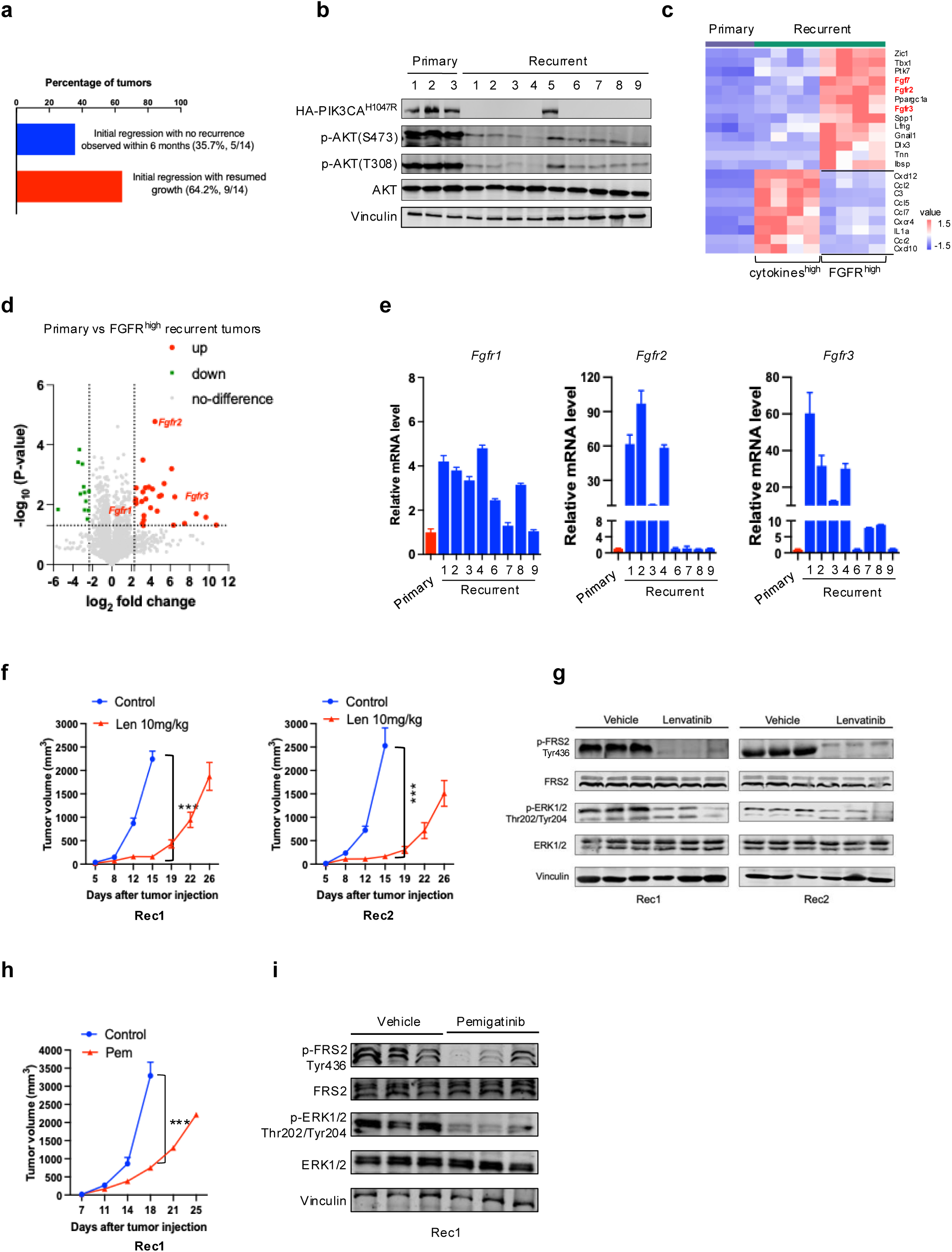
PIK3CA^H1047R^; Trp53^-/-^; Myc tumors recur after PIK3CA^H1047R^ withdrawal and exhibit FGFR-dependent proliferation. **(a)** Summary of primary tumor responses following doxycycline withdrawal: 5 out of 14 tumors (35.7%) regressed completely without regrowth (blue bar), whereas 9 out of 14 tumors (64.3%) initially regressed but subsequently regrew (red bar). **(b)** Immunoblot analysis showing expression levels of PIK3CA and associated downstream signaling pathways in primary and recurrent tumors. **(c)** Heat map illustrating significantly altered gene expression in recurrent tumors relative to primary tumors. **(d)** Volcano plot illustrating significantly altered gene expression in FGFR^high^ recurrent tumors. **(e)** FGFR transcript levels in primary versus recurrent tumors. **(f, h)** Tumor growth curves of recurrent tumors Rec1 and Rec2 treated with lenvatinib or pemigatinib (n=8-10 per group). **(g, i)** Immunoblot analysis showing reduced phosphorylation of FRS2 and ERK1/2 after 12 days of treatment.

Recurrent tumors preserved serous-like histopathological features (**Supplementary Fig. 2b and Supplementary Fig. 1e**). Functionally, recurrent tumors exhibited markedly reduced PI3K pathway activity and resistance to BYL719, with the except of Rec5, which retained PIK3CA-H1047R expression and remained sensitive to BYL719 (**Fig. 2b and Supplementary Fig. 2c, d**). These findings demonstrate that although PIK3CA-H1047R is required for tumor formation, most tumors eventually acquire PIK3CA-H1047R- independent mechanisms to sustain growth.

Transcriptomic profiling further revealed distinct gene expression patterns between primary and recurrent tumors (**Fig. 2c**). Recurrent tumors clustered into two major subgroups: one characterized by high expression of FGF/FGFR family members (FGF7, FGFR1, FGFR2, FGFR3), and another defined by elevated expression of multiple cytokines (**Fig. 2c-e and Supplementary Fig. 2e-g**). Quantitative PCR analysis confirmed elevated expressions of FGFR1, FGFR2, and FGFR3 in FGFR-high recurrent tumors (**Fig. 2e**). A similar pattern was observed in tumors that acquired resistance to BYL719, which showed increased FGFR expression at the time of resistance (**Supplementary Fig. 2h**).

FGFR signaling has been implicated in therapy resistance and cancer progression across multiple tumor types ^38, 39^. To explore the functional relevance of elevated FGFR expression in recurrent tumors, we transplanted FGFR-high recurrent tumors into wild-type FVB mice and treated them with either lenvatinib or pemigatinib, a more FGFR-selective inhibitor approved for FGFR-driven cancers ^40, 41^. Both agents effectively suppressed tumor growth and were accompanied by reduced phosphorylation of fibroblast growth factor receptor substrate 2 (FRS2) and extracellular signal-regulated kinase (ERK) (**Fig. 2f-i**). Collectively, these results indicate that a subset of tumors recurring after PIK3CA^H1047R^ withdrawal becomes dependent on FGFR signaling for sustained proliferation.

### FGFR-driven resistance and clonal dynamics in recurrent tumors after PIK3CA-H1047R inhibition

To decipher the molecular programs and clonal dynamics driving residual and recurrent disease after PIK3CA-H1047R inhibition, we performed single-nucleus RNA sequencing (snRNA-seq) on primary tumors and FGFR-high recurrent tumors (analyzing 23,116 cells with a median of ∼2,300 genes detected per cell) (**Supplementary Fig. 3a**). This analysis identified five major cell populations: epithelial cells, fibroblasts, endothelia cells, myeloid cells, and T cells (**Supplementary Fig. 3b, c**). RNA velocity analysis of epithelial cells further resolved four distinct epithelial clusters characterized by activation of unique signaling programs (**Fig. 3a, b**).

**Figure 3.**
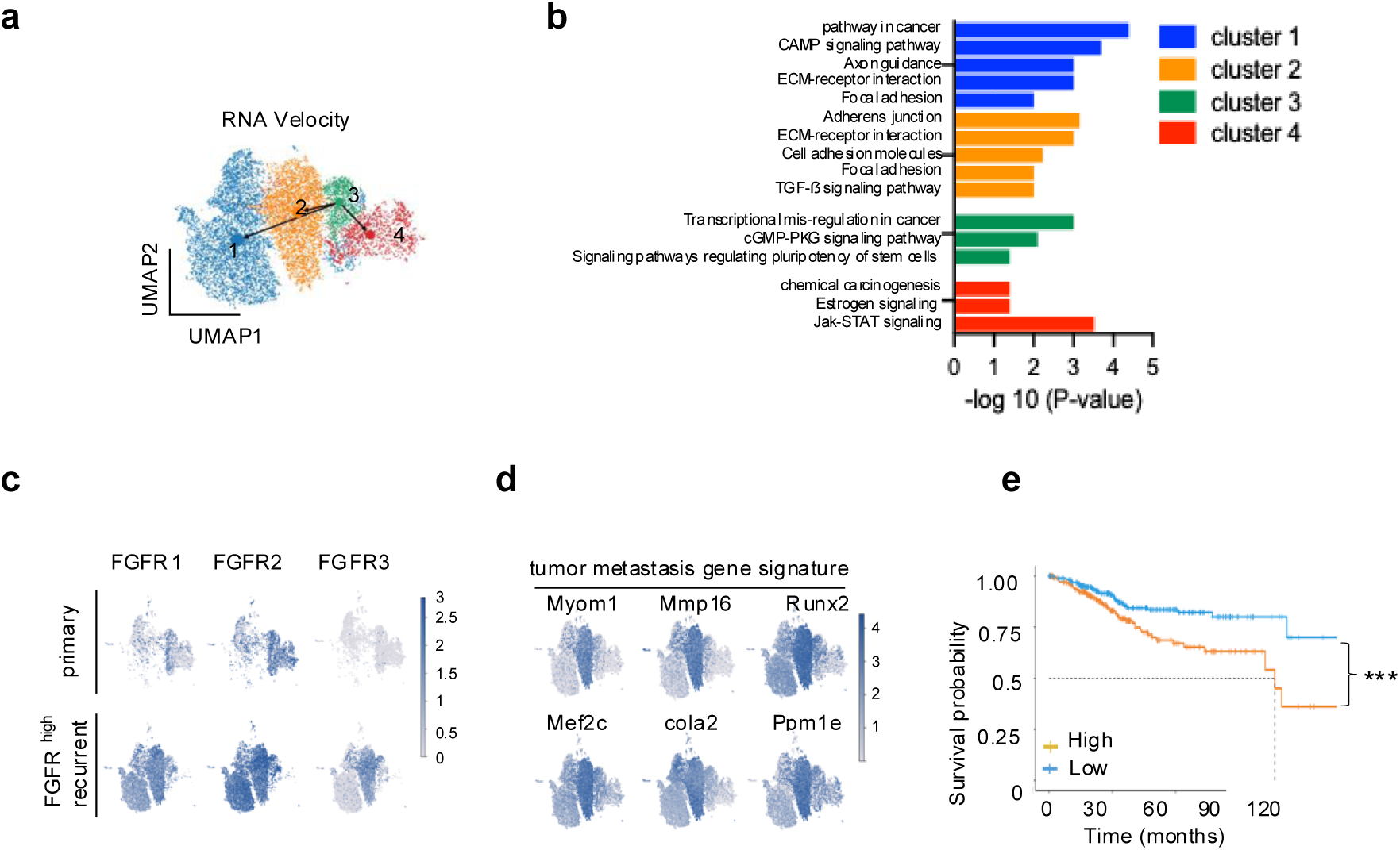
snRNA-seq reveals the evolutionary trajectory of FGFR^+^ tumor cells driving recurrence. **(a)** RNA velocity analysis overlaid on the UMAP projection suggests that FGFR-high recurrent tumor cells arise from epithelial cluster 3 present in primary tumors. **(b)** KEGG pathway enrichment analysis of differentially expressed genes (DEGs) in epithelial cells identified in the snRNA-seq dataset. **(c, d)** UMAP feature plots showing that FGFR1- and FGFR2-high tumor cells are already present in primary tumors, whereas FGFR3 expression emerges predominantly in recurrent tumors, indicating intrinsic versus acquired resistance programs. **(e)** Genes enriched in epithelial cluster 2, characterized by high FGFR1-3 expression, are associated with tumor metastasis signatures and reduced overall survival in TCGA UCEC patients. Survival analysis were performed using the R packages survival (version 3.5.5) and survminer (version 0.4.9).

Detailed examination of FGFR1-3 expression patterns provided important insights into resistance mechanisms. Tumor cells with high FGFR1 and FGFR2 expression were already present in the primary tumors (**Fig. 3c and Supplementary Fig. 2e, f**), consistent with an intrinsic or de novo resistance mechanism. In contrast, FGFR3 expression emerged predominantly in the recurrent tumors, indicative of an acquired resistance mechanism (**Fig. 3c and Supplementary Fig. 2e, f**). Notably, epithelial cluster 2 in recurrent tumors showed coordinated enrichment of FGFR1-3 expression together with TGF-β signaling and cell adhesion pathways commonly associated with metastasis progression (**Fig. 3a-d**).

To assess the clinical relevance of these transcriptional programs, we derived a gene signature enriched in epithelial cluster 2. High expression of this FGFR-enriched signature correlated with increased expression of metastasis-associated gene programs and significantly reduced overall survival in patient datasets (**Fig. 3e**). These findings suggest that FGFR-high tumor cell populations represent both a prognostic marker and a potential therapeutic vulnerability.

To evaluate microenvironmental remodeling during tumor recurrence, we next compared immune compartments between primary and recurrent tumors. FGFR-high recurrent tumors (Rec1) exhibited an marked increase in M2-type TAMs, accompanied by a reduction in M1-TAMs and total T-cell infiltration relative to primary tumors (APM) (**Supplementary Fig. 3d, e**). Within the T-cell compartment, we observe a pronounced depletion of cytotoxic CD8⁺ T cells and a relative enrichment of exhausted T cells and regulatory T cells (Tregs), populations associated with immune suppression (**Supplementary Fig. 3f**). Together, these data define a coordinated immune remodeling program in FGFR-high recurrent tumors, characterized by M2-dominant TAMs, loss of cytotoxic CD8⁺ T cells, and expansion of Tregs, which collectively promote immune escape and tumor recurrence.

### FGFR signaling mediates resistance to PIK3CA-targeted therapy

To further determine the role of FGFR signaling in resistance to PIK3CA-directed therapy, we treated a panel of ten PIK3CA-mutant human EC cell lines with the PI3Kα inhibitors BYL719 and GDC-0077 ^42, 43, 44, 45, 46, 47, 48, 49^. Higher FGFR expression correlated with reduced sensitivity to both inhibitors, as reflected by increased IC_50_ values (0.9-27 μM for BYL719 and 0.6-20 μM for GDC-0077), with the exception of ARK1, which harbors HER2 amplification ^23^ (**Fig. 4a and Supplementary Fig. 4a**).

**Figure 4.**
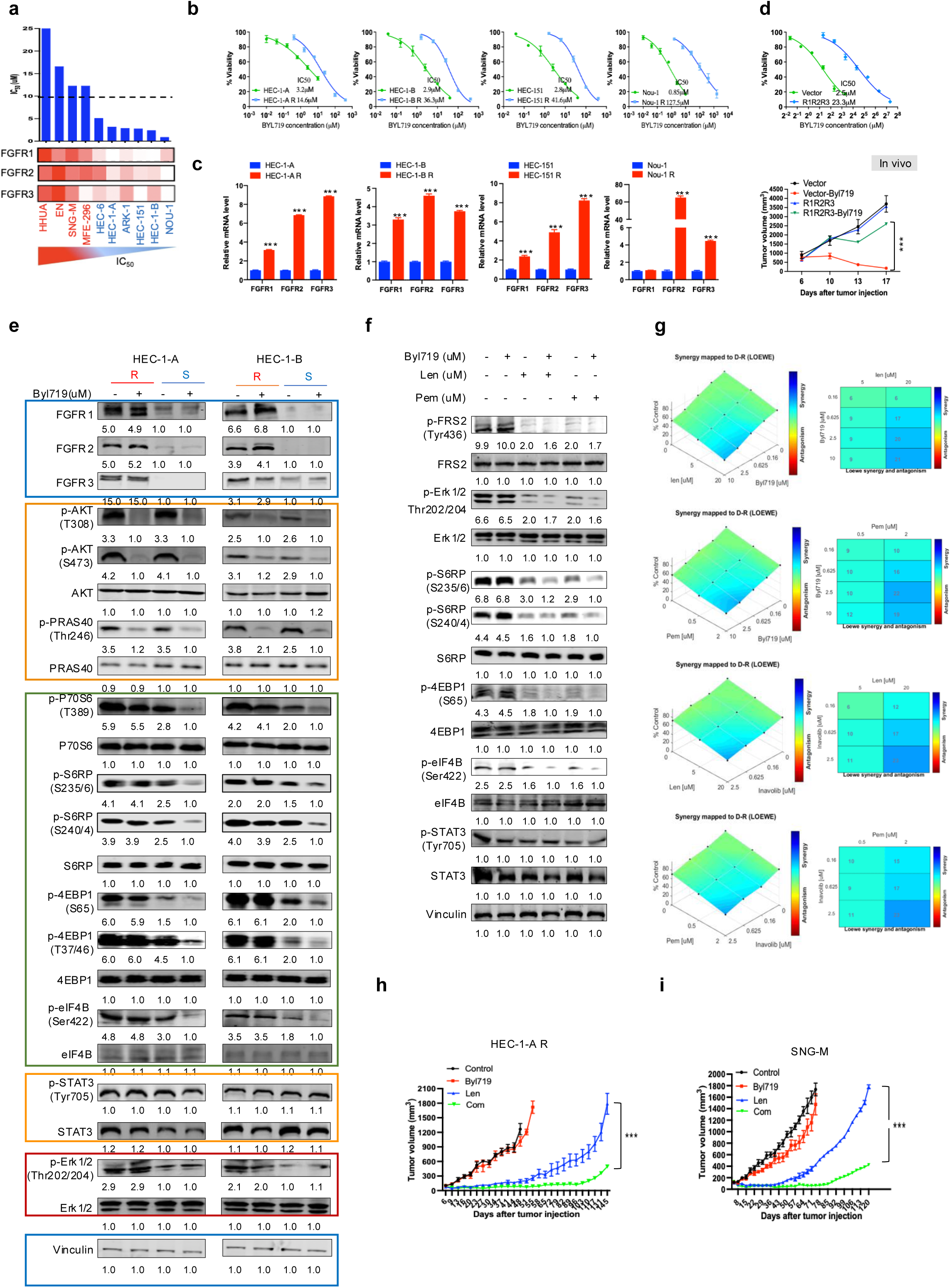
FGFR-driven MAPK-mTOR activation contributes to resistance to PI3Kα inhibition. **(a)** Correlation between FGFR1-3 expression levels and BYL719 IC_50_ values across ten human EC cell lines harboring PIK3CA mutation. **(b)** Dose-response analyses comparison proliferation of four BYL719-sensitive cell lines and their matched resistant counterparts following treatment with increasing doses of BYL719. **(c)** Relative FGFR1-3 expression levels in BYL719-sensitive cell lines compared with their resistant derivatives. **(d)** BYL719 sensitivity (IC_50_, top) and in vivo tumor growth (bottom) of primary tumor cells stably overexpressing FGFR1-3 or carrying an empty vector following BYL719 treatment (n = 8-10 per group). **(e)** Immunoblot analysis of PI3K/AKT, MAPK, and STAT3 signaling in BYL719-sensitive (S) and - resistant (R) cells treated for 24 hours with 1 μM BYL719. **(f)** Immunoblot analysis of BYL719-resistant cells treated with BYL719 (1 μM), lenvatinib (20 μM), pemigatinib (2 μM), or the indicated combinations, showing suppression of MAPK-mTOR signaling upon FGFR inhibition. **(g)** Isobologram plots showing additive or synergistic effects of BYL719 or inavolisib combined with Lenvatinib or pemigatinib on HEC-1-A R cell viability, analyzed using the Combenefit synergy-antagonism model. **(h, i)** Tumor growth curves of xenografts derived from BYL719-resistant HEC-1-A R (h) and intrinsically resistant SNG-M tumors (i) treated with the indicated agents (n = 6-10 per group).

To directly test whether FGFR contributes to acquired resistance, we selected four EC cell lines (HEC-1-A, HEC-1-B, HEC-151, and Nou-1) with low baseline FGFR expression and initial sensitivity to BYL719. Chronic exposure to increasing concentrations of BYL719 generated resistant derivatives, using 10 μM as the resistance threshold based on clinically achievable plasma levels ^30^. All resistant lines displayed IC_50_ values greater than 10 μM and exhibited marked upregulation of FGFR1-3 compared with their parental counterparts (**Fig. 4b, c**). Similar results were obtained with GDC-0077 in HEC-1-A iR cells, an inavolisib-resistant derivative generated by stepwise dose escalation (**Supplementary Fig. 4b**). In addition, enforced expression of FGFR1-3 in primary APM tumor cells was sufficient to confer resistance to BYL719 (**Fig. 4d**).

Detailed signaling analyses revealed that although BYL719 effectively suppressed phosphorylation of AKT and its downstream target PRAS40 in both sensitive and resistant cells, phosphorylation of downstream mTOR effectors, including S6, eIF4B, and 4EBP1, persisted in BYL719-resistant HEC-1-A R and HEC-1-B R cells (**Fig. 4e**). Co-treatment with a PI3Kα inhibitor (BYL719 or GDC-0077) together with an FGFR inhibitor (lenvatinib or pemigatinib) effectively abolished phosphorylation of S6, eIF4B, and 4EBP1, leading to a marked reduction in proliferation in BYL719-resistant HEC-1-A R and SNG-M cells (**Fig. 4f, g and Supplementary Fig. 4c**). Similar effects were observed upon FGFR knockdown, which resensitized SNG-M and HEC-1-A R to BYL719 (**Supplementary Fig. 4d**).

Consistent with these findings, FGFR-high recurrent Rec1 tumors exhibited marked upregulation of FGFR1-3, accompanied by increased ERK1/2 phosphorylation and activation of mTOR downstream effectors (p-S6RP and p-4EBP1), relative to primary APM tumors, whereas STAT3 phosphorylation remained largely unchanged (**Supplementary Fig. 4e**). shRNA-mediated knockdown of FGFR1-3 in Rec1 tumors substantially reduced p-ERK1/2 and p-S6RP/p-4EBP1 levels, indicating that FGFR upregulation in the recurrent setting predominantly sustains MAPK-mTOR signaling (**Supplementary Fig. 4e**). Notably, AKT phosphorylation was robust in primary APM tumors but was not maintained in Rec1 tumors, consistent with loss of PIK3CA^H1047R^ activity. Thus, AKT is not the dominant downstream effector of FGFR-driven signaling in this model, reflecting an uncoupling of mTOR output from PI3K-AKT signaling in the resistant state. Accordingly, FGFR blockade in this context primarily suppresses the FRS2-MAPK-mTOR axis, with minimal effects on STAT3 and PI3K-AKT signaling (**Supplementary Fig. 4f**).

We next validated these findings in vivo. BYL719-resistant HEC-1-A R xenografts exhibited a markedly enhanced response to combined BYL719 and lenvatinib treatment compared with lenvatinib monotherapy (**Fig. 4h and Supplementary Fig. 4g**). This combination also demonstrated strong efficacy in intrinsically BYL719-resistant SNG-M xenografts (**Fig. 4i and Supplementary Fig. 4h, i**), with similar effects observed using the FGFR-selective inhibitor pemigatinib in combination with BYL719 (**Supplementary Fig. 4j**), indicating its potential to overcome both acquired and intrinsic resistance to PI3Kα inhibition. Similarly, in the APM model rendered resistant after prolonged BYL719 exposure (termed APM-R), FGFR inhibition restored sensitivity to PI3Kα blockade (**Supplementary Fig. 4k**). In contrast, in BYL719-sensitive HEC-1-A tumors, the combination produced a synergistic but less pronounced effect (**Supplementary Fig. 4l, m**). Collectively, these data indicate that PI3Kα and FGFR co-inhibition is at least additive and frequently synergistic across both PI3Kα-sensitive and - resistant settings, with synergy most evident in the resistant, FGFR-driven context.

### FGFR overexpression promotes tumor immune evasion in EC

Analysis of TCGA EC datasets revealed a significant inverse correlation between FGFR expression and CD3D levels, indicating reduced T-cell infiltration in FGFR-high tumors (**Fig. 5a**). Elevated FGFR2/3 expression was preferentially enriched in uterine serous carcinoma (**Fig. 5a**). Moreover, FGFR- amplified tumors (n=51) exhibited significantly lower expression of major histocompatibility complex class I (MHC-I) genes (HLA-A, HLA-B, and HLA-C) compared with non-amplified tumors (n=495) (**Fig. 5b**), consistent with impaired antigen presentation. CIBERSORT-based immune deconvolution further demonstrated that FGFR-high tumors contained fewer CD8⁺ T cells but a higher proportion of M2 macrophages, supporting a more immunosuppressive tumor microenvironment associated with FGFR activation (**Supplementary Fig. 5a**). Although changes in some immune subsets were less pronounced, FGFR-high tumors showed a general shift toward a less inflamed phenotype, with reduced effector T-cell signatures and a myeloid compartment skewed toward M2/TAM-like populations (**Supplementary Fig. 5a**).

**Figure 5.**
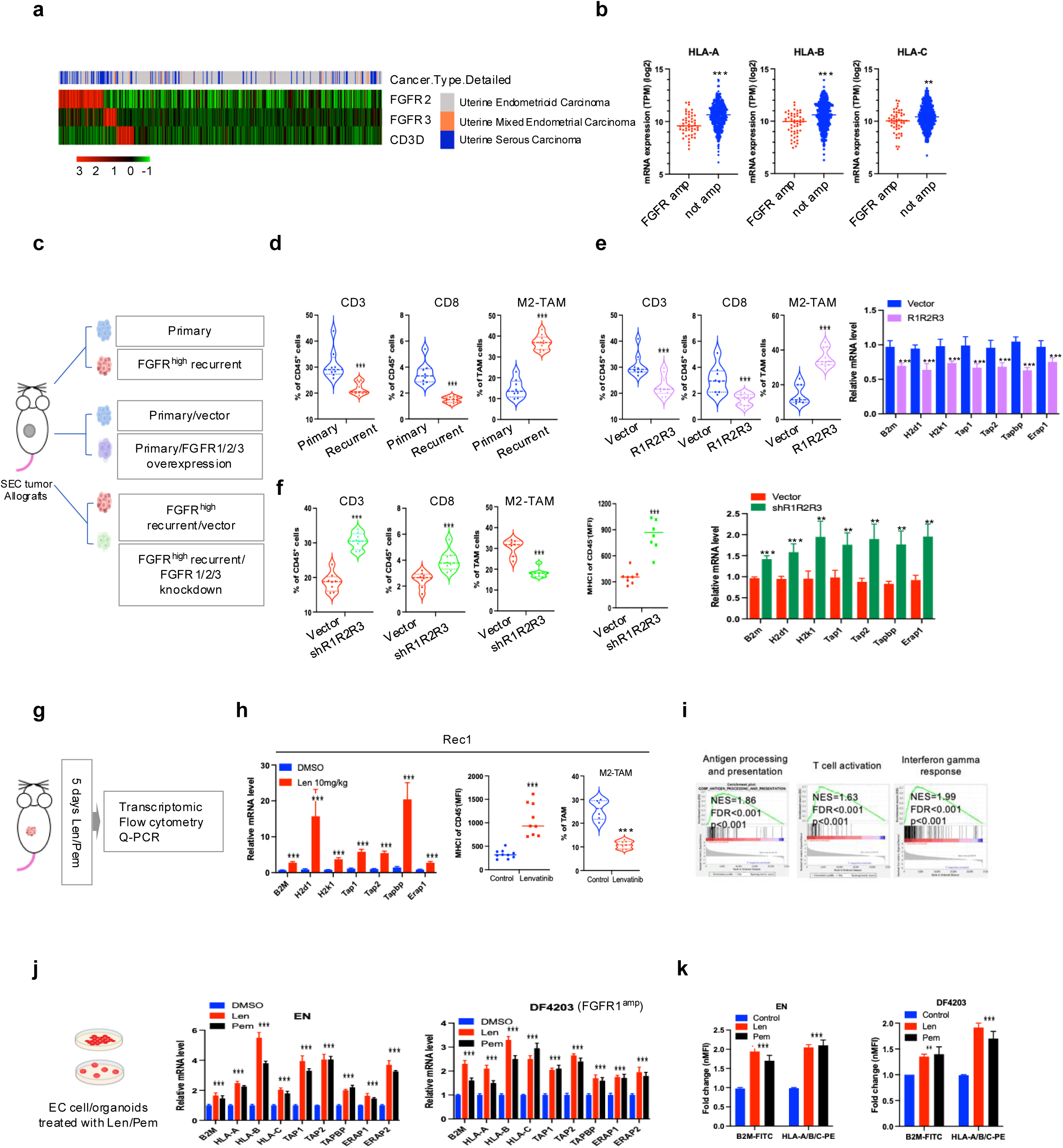
FGFR inhibition enhances antigen presentation and reduces M2-like tumor-associated macrophages. **(a)** TCGA analysis revealed an inverse relationship between FGFR expression and CD3D levels (https://www.cbioportal.org/study/summary?id=ucec_tcga_pan_can_atlas_2018). The cohort includes 397 uterine endometrioid carcinomas, 109 uterine serous carcinomas, and 21 mixed endometrioid carcinomas. Heatmap and oncoprint visualizations were generated using the R package pheatmap to display FGFR and CD3D expression (log_2_ odds ratio <-2). **(b)** Differential gene expression analysis comparing FGFR-amplified (n=51) and non-amplified (n=495) TCGA tumors (https://www.cbioportal.org/study?id=ucec_tcga_gdc). **(c)** Schematic of the in vivo experimental design. **(d-f)** Flow cytometry and RT-qPCR analyses of CD3^+^ and CD8^+^ T cells, M2-TAMs, and antigen-presentation gene expression in primary tumors, recurrent tumors, primary tumors overexpressing FGFR1/2/3, and recurrent tumors with FGFR knockdown (n=8-10 per group). **(g-i)** GSEA, RT-qPCR, and flow cytometry analyses demonstrating upregulation of antigen presentation-related genes and a reduction in M2-TAMs in FGFR-high recurrent Rec1 tumors following short-term lenvatinib treatment. **(j-k)** Expression of antigen-presentation genes in EC cell lines and PDOs after 7 days of FGFR inhibitor treatment. PDO DF4203 harbors FGFR1 amplification.

Importantly, this immune landscape was recapitulated in our preclinical models (**Fig. 5c**). Both recurrent Rec1 tumors with spontaneous FGFR overexpression (**Fig. 5d**) and primary tumors engineered to overexpress FGFRs (**Fig. 5e and Supplementary Fig. 5b**) exhibited reduced T-cell infiltration, increased M2-type TAMs, and diminished MHC-I-mediated antigen processing and presentation. In contrast, tumors with shRNA-mediated FGFR knockdown displayed increased CD3⁺ and CD8⁺ T-cell infiltration, reduced M2 macrophages abundance, and enhanced MHC-I expression (**Fig. 5f**). Together, these findings indicate that elevated FGFR signaling drives an immunosuppressive tumor microenvironment in both human EC and the APM model, aligning the immune features of our preclinical system closely with those observed in patient tumors.

We next tested whether FGFR inhibition could reverse this immunosuppressive state. Using a FGFR-high serous-like EC allograft model, we performed immune and transcriptomic profiling following short-term (5 days) FGFR inhibitor treatment (**Fig. 5g**). FGFR blockade significantly increased the expression of MHC-I- associated and antigen-presentation genes and reduced M2-TAM populations (**Fig. 5h and Supplementary Fig. 5c**). GSEA of 4,604 cancer-related genes revealed strong enrichment of antigen processing and presentation pathways in FGFR inhibitor-treated tumors (**Fig. 5i**), a pattern that was reproduced in additional FGFR-high recurrent Rec2 tumors (**Supplementary Fig. 5d**). Concordantly, human EC cell lines and patient-derived organoids (PDOs) harboring FGFR overexpression or amplification also exhibited enhanced antigen presentation following lenvatinib or pemigatinib treatment (**Fig. 5j, k**).

Collectively, these results support a model in which the primary immunomodulatory effects of lenvatinib in our system are mediated through tumor-cell-intrinsic FGFR blockade, rather than direct suppression of immune cells. Consistent with this, treatment of naïve, tumor-free mice with lenvatinib for 12 days did not alter the frequencies of M2-like macrophages or T cells in the spleen or peripheral blood (**Supplementary Fig. 5e**), arguing against a major direct systemic immune cell-intrinsic effect of the drug in the absence of tumor.

To define the downstream signaling mechanisms underlying FGFR-driven immune suppression, we next interrogated major pathways activated downstream of FGFR, including MAPK, STAT3, and PI3K-AKT signaling. Pharmacologic MAPK inhibition most closely recapitulated the immunomodulatory effects of FGFR blockade, increasing MHC-I expression on tumor cells both in vitro and in vivo Rec1 tumor allografts, while concomitantly reducing M2-type TAMs infiltration (**Supplementary Fig. 5 f, i**). In contrast, inhibition of STAT3 or PI3K did not produce comparable changes in antigen-presentation gene expression (**Supplementary Fig. 5 g, h**). Moreover, combined inhibition of MAPK and PI3K signaling markedly restored BYL719 sensitivity in PI3Kα-resistant APM-R tumors, which were generated by long-term BYL719 dose escalation and characterized by FGFR overexpression (**Supplementary Fig. 2h**). This combination led to robust tumor growth suppression in vivo (**Supplementary Fig. 5j**). Notably, mice maintained stable body weight during combined MAPK inhibitor and BYL719 treatment, indicating good tolerability (**Supplementary Fig. 5j**). Together, these findings identify MAPK-ERK signaling as a key downstream mediator of FGFR-driven immune suppression and therapeutic resistance, providing a mechanistic rationale for combination strategies integrating PI3Kα and MAPK pathway inhibition alongside FGFR-targeted therapy.

### FGFR inhibition enhances antitumor immunity and augments immunotherapy

The observed increase in tumor antigen presentation and enhancement of antitumor T cell response suggested that combining FGFR inhibition with PD-1 blockade might further improve therapeutic efficacy. To test this hypothesis, we treated mice bearing FGFR-high recurrent tumors with lenvatinib alone or in combination with an anti-PD-1 antibody (**Fig. 6a**). Lenvatinib monotherapy produced transient tumor control followed by regrowth, whereas the combination led to markedly stronger and more durable tumor suppression (approximately 85% reduction in tumor volume by day 27; **Fig. 6b**), with comparable effects observed in additional FGFR-overexpressing recurrent tumors and in models treated with other FGFR inhibitors (**Supplementary Fig. 6a, b**).

**Figure 6.**
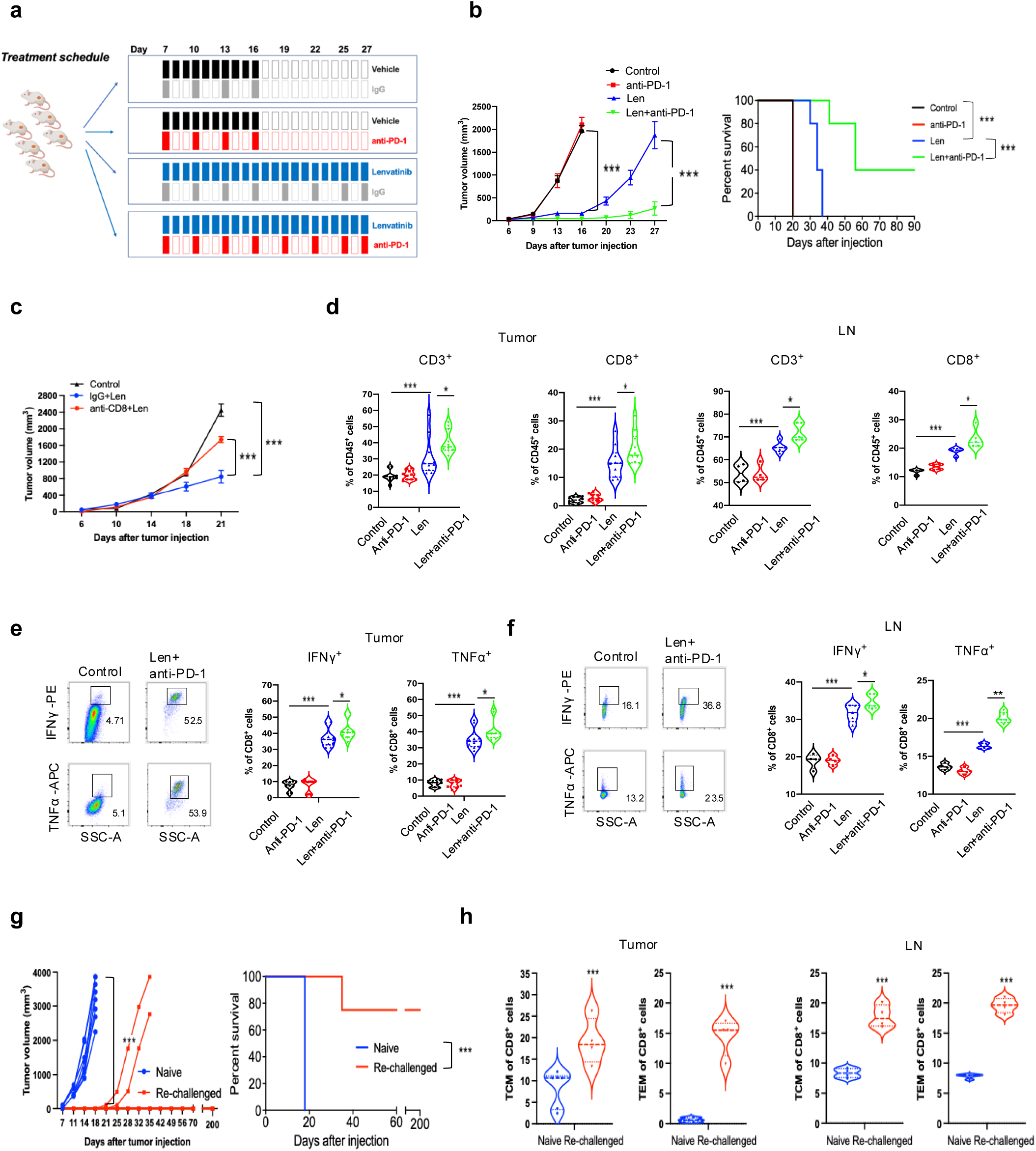
FGFR blockade enhances T-cell-mediated antitumor immunity and augments the therapeutic efficacy of PD-1 blockade. **(a)** Schematic of the experimental design for treating recurrent tumor allografts with the indicated agents for 21 days. **(b)** Tumor volume curves and Kaplan-Meier survival analysis of recurrent Rec1 tumors treated with lenvatinib alone or in combination with anti-PD-1 therapy (n = 8-10 per group). **(c)** Tumor growth curves of mice treated with lenvatinib in the presence or absence of an anti-CD8 neutralizing antibody (n=6 per group), indicating a requirement for CD8⁺ T cells in lenvatinib-mediated tumor control. **(d-f)** Flow cytometric analysis of intratumoral and draining lymph node (dLNs) CD3^+^ and CD8^+^ T-cell populations and effector cytokine production following treatment with the indicated agents (n=8-10 per group). **(g)** Tumor growth curves (n = 10 per group) and survival analysis (n = 5 per group) of recurrent tumors transplanted into Naïve mice or mice rechallenged after complete tumor regression. **(h)** Six days after recurrent tumor injection into Naïve or rechallenged mice, tumors and dLNs were harvested for memory T cell analysis. Flow cytometry was used to quantify CD8 TEM (CD45^+^ CD3^+^ CD8^+^ FOXP3^-^ CD62L^-^ CD44^+^ KLRG1^+^) and CD8 TCM (CD45^+^ CD3^+^ CD8^+^ FOXP3^-^ CD62L^+^ CD44^+^ KLRG1^-^).

Depletion of CD8^+^ T cells in immunocompetent FVB mice substantially attenuated the anti-tumor activity of lenvatinib (**Fig. 6c and Supplementary Fig. 6c**), indicating that CD8⁺ T cells are required for lenvatinib-mediated tumor control. Immunological profiling further revealed that lenvatinib monotherapy increased the production of effector cytokines (e.g., IFN-γ and TNF-α) by CD8^+^ T cells, an effect that was further enhanced by combination with anti-PD-1 therapy (**Fig. 6d-f and Supplementary Fig. 6d, e**), with similar results observed in tumors treated with other FGFR inhibitors (**Supplementary Fig. 6f, g**).

Notably, combined lenvatinib and anti-PD-1 therapy achieved complete tumor regression in ∼40% of mice (**Fig. 6b**). To assess long-term immunity, mice that achieved complete responses were re-challenged with tumor cells four weeks after treatment cessation. Whereas all naive control mice developed tumors within one month, 80% of re-challenged mice remained tumor-free (**Fig. 6g**), consistent with the establishment of durable immune protection. Mechanistically, analysis performed six days after tumor re-inoculation revealed significantly increased frequencies of effector memory (TEM) and central memory (TCM) T cells in both tumors and draining lymph nodes compared with naive mice (**Fig. 6h**). Together, these findings demonstrate that FGFR inhibition combined with PD-1 blockade not only enhances immediate antitumor immunity but also induces durable immunological memory capable of preventing tumor recurrence.

## DISCUSSION

SEC remains one of the most aggressive EC subtypes, yet progress in understanding its molecular pathogenesis and therapeutic vulnerabilities has been hindered by the lack of in vivo models that faithfully reproduce its defining biological features. SEC is characterized by extensive copy-number alterations, genomic instability, hallmark serous histology, and distinctive immune interactions, attributes largely absent from existing endometrioid-based models ^50^. To address this gap, we generated a clinically relevant, immunocompetent SEC-like mouse model – termed APM - by integrating oncogenic PIK3CA mutation, Trp53 loss, and Myc overexpression, three genetic alterations that collectively define high-risk serous tumors^4^. This APM model system faithfully recapitulates key molecular hallmarks of human SEC, including a CNV-high genomic profile, enrichment of G2M checkpoint activation, replication stress programs, E2F and MYC transcriptional signatures, and classic serous morphology with high-grade architectural and nuclear atypia. Because APM tumors and their derived recurrent tumors evolve under selective pressures imposed by genomic instability, replication-associated stress, and immune surveillance, this system provides a unique opportunity to interrogate therapeutic pressure-driven clonal evolution while preserving intact tumor-immune interactions.

By integrating this model with analyses of EC cell lines, xenografts, and patient datasets, we identified FGFR signaling as a central determinant of resistance to PI3Kα-targeted therapy. Single-cell transcriptomic profiling reveals a functional hierarchy within the FGFR family: FGFR1 and FGFR2 are predominantly associated with intrinsic resistance, whereas FGFR3 becomes selectively enriched during acquired resistance, consistent with adaptive signaling rewiring under therapeutic pressure. Mechanistically, FGFR activation sustains mTORC1 activity and enhances MAPK signaling despite effective PI3Kα blockade, as evidenced by persistent phosphorylation of S6 and 4EBP1 together with increased ERK phosphorylation, while AKT and STAT3 activity remain largely unchanged. These findings position persistent MAPK-mTORC1 signaling downstream of FGFR as the dominant proliferative driver enabling escape from PI3Kα inhibition. Accordingly, dual PI3Kα and FGFR inhibition produced markedly enhanced antitumor effects in vivo, validating FGFR activation as a therapeutically actionable resistance node.

These mechanistic observations align with extensive preclinical evidence demonstrating that FGFR-mediateed pathway rewiring enables tumor cells to escape PI3K or receptor tyrosine kinase (RTK) inhibition ^39, 51, 52, 53^, and that dual PI3K-FGFR blockade yields robust synergy across multiple cancer types ^54, 55^. Early-phase clinical trials evaluating alpelisib in combination with the FGFR1-3 inhibitor infigratinib further underscore the translational potential of this strategy, although overlapping toxicities, such as hyperglycemia and hyperphosphatemia, emphasize the importance of optimized dosing regimens and biomarker-guided patient selection ^56^. Together with our mechanistic data, these findings provide strong biological and translational justification for pursuing PI3K-FGFR combination therapies in molecularly defined subsets of SEC.

Beyond tumor-intrinsic signaling, our data demonstrate that FGFR activation also orchestrates immune evasion. Elevated FGFR expression suppresses tumor immunogenicity by downregulating MHC-I/HLA-mediated antigen processing and presentation and by promoting an immunosuppressive tumor microenvironment characterized by increased infiltration of M2-type TAMs. Tumor-cell-specific FGFR depletion phenocopied the immunologic impact of systemic FGFR inhibition, confirming that these effects arise from tumor-intrinsic FGFR signaling rather than off-target effects on immune cells. Importantly, FGFR inhibition restored antigen presentation, enhanced CD8⁺ T-cell activation, improved responsiveness to PD-1 blockade, and promoted durable immunologic memory. These findings support a conceptual model in which FGFR signaling functions as a lineage-specific immune checkpoint that lowers tumor antigenicity and conditions an immunosuppressive microenvironment.

Our observations integrate into a broader mechanistic and clinical framework supporting FGFR-immune checkpoint blockade (ICB) combination strategies. In FGFR3-mutant bladder cancer, FGFR inhibition enhances anti-PD-1 efficacy by increasing CD8⁺ T-cell infiltration and restraining regulatory T-cell expansion ^57^. Parallel immune-activating effects of FGFR blockade have been reported in hepatocellular, gastric, and lung cancers, where FGFR inhibition increases IFNγ production, boosts MHC-I expression, and strengthens T-cell activation ^58, 59^. Clinically, the combination of lenvatinib and pembrolizumab has produced substantial and clinically meaningful response rates and survival benefits in advanced EC ^14^, while early-phase trials combining the FGFR4 inhibitor FGF401 with spartalizumab have demonstrated manageable safety profiles and preliminary efficacy across solid tumors ^60^.

However, FGFR pathway activation alone is not sufficient to uniformly predict benefit from FGFR-targeted immunotherapy. Although lenvatinib plus PD-1 blockade has shown substantial activity, responses remain highly heterogeneous, and FGFR-high status has not consistently emerged as a predictive biomarker. Our study provides mechanistic insight into this heterogeneity by revealing the emergence of two distinct recurrent tumor states following PIK3CA withdrawal. One state is characterized by FGFR upregulation coupled to immune suppression mediated through impaired antigen presentation and macrophage polarization. Tumors within this state derive substantial benefit from combined FGFR inhibition and PD-1 blockade, consistent with FGFR functioning as a lineage-specific immune checkpoint in this context.

In contrast, we identified a subset of recurrent tumors characterized by increased cytokine expression in the absence of FGFR amplification or overexpression. These tumors exhibit features of an FGFR-independent immune-resistant program and appear to evade immune surveillance through inflammation-linked remodeling of the tumor microenvironment. Sustained cytokine signaling has been associated with immune dysfunction, including impaired antigen presentation, cytokine-driven myeloid cell infiltration, and the development of T-cell exhaustion, collectively promoting adaptive immune resistance ^61, 62, 63^. Accordingly, cytokine-high recurrent tumors may be less amenable to immunotherapeutic strategies that rely on reinvigoration of cytotoxic T-cell responses, highlighting a potential limitation of immune checkpoint blockade in this subset. This alternative resistance state is not explained by FGFR-mediated immune suppression and provides a mechanistic framework for the observed heterogeneity in responses to FGFR-immune checkpoint blockade combinations, underscoring the need for biomarker frameworks that integrate inflammatory programs in addition to FGFR activity.

In conclusion, our study establishes a dual role for FGFR signaling in mediating both therapy resistance and immune evasion in SEC. By demonstrating that FGFR inhibition augments the efficacy of both PI3Kα-targeted therapy and immune checkpoint blockade, we provide a compelling rationale for therapeutically integrating FGFR blockade into combination regimens (**Supplementary Fig. 6h**). At the same time, overlapping toxicities, particularly in PI3K-FGFR combinations, remain a major clinical challenge, requiring careful dose optimization, scheduling, and supportive management. Ultimately, realizing the full clinical potential of these approaches will depend on biomarker-guided patient stratification, precision dosing paradigms, and rational combination designs that maximize antitumor potency while maintaining tolerability.

## Methods

### Mice

All animal experiments were performed in accordance with protocols (03-111, 06-034, 02-127) approved by the Dana-Farber Cancer Institute (DFCI) Institutional Animal Care and Use Committee (IACUC). The TetO-PIK3CA^H1047R^ transgenic mouse line was maintained in our laboratory as previously described ^33^. The Tp53^loxP/loxP^ mouse line (FVB.129P2-Trp53tm1Brn/Nci, #01XC2) was obtained from the National Cancer Institute Mouse Repository. Both strains were backcrossed for more than ten generations onto the FVB/N background before being intercrossed to generate homozygous lines. FVB/NJ and athymic nude mice were purchased from The Jackson Laboratory.

### Cell culture

The following human EC cell lines were used in this study: HEC-1-A and HEC-1-B (ATCC); MFE-296 (Sigma Aldrich); ARK1 (courtesy of Dr. Dipanjan Chowdhury, Dana-farber Cancer Institute); HEC-151, HEC-6, and HHUA (kindly provided by Dr. Jessie A. Sanai, Memorial Sloan Kettering Cancer Center); and EN, NOU-1, and SNG-M (generously provided by Dr. Gordon B. Mills, MD Anderson Cancer Center). All cell lines were routinely tested for mycoplasma contamination and authenticated by short tandem repeat (STR) profiling.

Primary mouse tumor-derived cells were established from freshly isolated primary or recurrent uterine tumors. Tumors were minced and enzymatically digested to generate single-cell suspensions, which were cultivated in serum-free F-Medium (Ham’s F-12:DMEM, 1:3) supplemented with 25 ng/mL hydrocortisone, 5 μg/mL insulin, 8.5 ng/mL cholera toxin, 0.125 ng/mL epidermal growth factor (EGF), 100 μg/mL penicillin-streptomycin, and 5 μM Y-27632 (ROCK1 inhibitor). Following initial tumor cell selection, cultures were maintained in F-medium supplemented with 1% fetal bovine serum (FBS).

### Generation of PI3Kα inhibitor-resistant cell lines

To generate BYL719-resistant cell lines, HEC-1-A, HEC-1-B, HEC-151, and NOU-1 cells were continuously exposed to increasing concentrations of BYL719, starting at 1ug/ml. Cells were serially passaged with stepwise dose escalation until resistant populations proliferated at rates comparable to parental controls in the presence of 50 μg/mL BYL719. This process required approximately six months and yielded stable BYL719-resistant derivatives.

Similarly, a GDC-0077-resistant HEC-1-A line (HEC-1-A iR) was generated through gradual dose escalation of GDC-0077. Acquisition of resistance was confirmed by drug withdrawal for 48 hours, followed by comparison of dose-response profiles between resistant and parental cells, demonstrating markedly reduced sensitivity to the corresponding PI3Kα inhibitor.

### Tumor growth and treatment

Primary and recurrent tumor cells (5 × 10^5^- 2x10^6^) for orthotopic implantation were suspended in serum-free DMEM mixed with 40% Matrigel and injected into 6-8-week-old female mice. For tumor-infiltrating immune cell profiling, 5×10^6^ recurrent tumor cells were implanted. Tumor size was measured 2-3 times per week, and the tumor volume was calculated using a modified ellipsoid formula (0.5 × length × width^2^). Mice bearing APM tumors received doxycycline-containing chow (2000 ppm) throughout the experiment.

For therapeutic studies, BYL719 and lenvatinib were dissolved in 0.5% methylcellulose and administered orally once daily at doses of 20 mg/kg and 10 mg/kg, respectively. Pemigatinib and trametinib were dissolved in 10% DMSO/90% corn oil and administered orally once daily at 1 mg/kg. GDC-0077 was formulated in 10% DMSO, 40% PEG300, 5% Tween-80, and 45% saline, and administered by oral gavage once daily at 50 mg/kg. Anti-PD-1 antibody (clone, 332.8H3, mouse IgG1) was diluted in PBS and administered intraperitoneally at 200 μg per mouse every 3 days ^64^. Control IgG (RTK2758, Biolegend) and anti-CD8 antibody (clone 53.6.7, Biolegend) were diluted in PBS and administered intravenously at 200 μg per mouse once weekly. Animals were monitored daily for clinical signs of toxicity, including activity, appetite, grooming, hydration status, and body weight.

### Bioluminescence Imaging

Mice were administered D-luciferin (Gold Biotechnology) via intraperitoneal injection at a dose of 120 mg/kg. Ten minutes after injection, bioluminescent signals were acquired using an IVIS Spectrum imaging system (PerkinElmer). Images were processed and quantified using Living Image software (PerkinElmer).

### Tissue digestion

To generate single-cell suspensions, tumors were excised, minced and enzymatically dissociated in a collagenase/hyaluronidase buffer (DMEM supplemented with 5% FBS, 10 mM HEPES, 100 μg/mL penicillin-streptomycin, 20 μg/mL DNase I, and 1× collagenase/hyaluronidase). Tissues were incubated at 37 °C for 40-60 minutes with gentle agitation. Following digestion, samples were treated with ammonium-chloride-potassium (ACK) lysis buffer to remove red blood cells (RBCs) and then filtered through a 70 μm cell strainer to eliminate debris and undigested material.

Tumor-draining lymph nodes (TDLNs) were mechanically dissociated by gently passing the tissues through a 70 μm cell strainer using the plunger of a 5 mL syringe, followed by ACK lysis as described above.

### Cell Viability Assay

Tumor cells were seeded in 96-well plates at a density of 2500 cells per well and allowed to adhere overnight. Cells were then treated with the indicated drugs at varying concentrations for 72 hours. Cell viability was assessed using the CellTiter-Glo 2.0 assay (Promega, #G9242) according to the manufacturer’s instructions. Luminescence was measured and normalized to untreated controls, which were defined as 100% viability. Growth inhibition was calculated based on relative luminescence values. Dose-response curves were generated using nonlinear regression analysis in GraphPad Prism 8.2, and IC_50_ values were derived from the fitted curves.

### Generation of FGFR-deficient and FGFR-overexpressing cells

Five independent shRNA constructs targeting human FGFR1, FGFR2, and FGFR3, as well as five constructs targeting mouse Fgfr1, Fgfr2, and Fgfr3, were obtained from the Sigma Mission shRNA library. All shRNA candidates were screened across multiple tumor cell lines to evaluate knockdown efficiency. For each FGFR gene, the shRNA achieving the most robust and reproducible knockdown was selected for subsequent studies: human FGFR1 (TRCN0000121106), FGFR2 (TRCN0000000373), FGFR3 (TRCN0000231051); mouse Fgfr1 (TRCN0000023295), Fgfr2 (TRCN0000023715), and Fgfr3 (TRCN0000361155).

Lentiviral particles containing the selected shRNAs were produced and used to infect target cells, followed by selection with puromycin (1 μg/mL) to generate stable knockdown cell lines. Knockdown efficiency was confirmed by RT-qPCR and Western blotting analysis.

To generate FGFR-overexpressing cells, full-length coding sequences of Fgfr1, Fgfr2, and Fgfr3 were amplified and individually cloned into the pCDH-CMV-EF1-Puro lentiviral expression vector. Lentiviral supernatants were produced and used to transduce tumor cells, and stable integrants were selected with puromycin (1 µg/ml). Overexpression levels were validated using RT-qPCR and Western blotting.

### Immunohistochemistry

Tumor specimens were fixed overnight in 10% neutral-buffered formalin and subsequently transferred to 70% ethanol. Tissue processing, paraffin embedding, sectioning, and hematoxylin and eosin (H&E) staining were performed by the Harvard Rodent Histopathology Core. Histological evaluation of primary and recurrent tumors was conducted by independent pathologists at Harvard Medical School.

Immunohistochemical (IHC) was performed using standard protocols. The following primary antibodies were used: anti-HA (1:400, Cell Signaling Technology, #3724), anti-pan-cytokeratin (1:200, Abcam, #ab7753), anti-ERα (1:200, Thermo Fisher Scientific, #RM9101S0), anti-Ki67 (1:500, Abcam, #ab15580), anti-cleaved caspase-3 (1:400, Cell Signaling Technology, #9661), anti-FGFR1 antibody (1:50, Abcam, #ab76464), anti-FGFR2 antibody (1:50, Abcam, #ab106648), and anti-FGFR3 antibody (1:50, Abcam, #ab180906).

### RNA extraction and reverse transcription-quantitative PCR (RT-qPCR)

Total RNA was isolated using the RNeasy Plus Mini Kit (QIAGEN, #74134) according to the manufacturer’s instructions. cDNA was synthesized from 2 µg of RNA using the iScript Reverse Transcription Supermix (Bio-Rad, #1708841). Quantitative PCR was performed using a real-time PCR system following standard protocols. Gene expression levels were normalized to endogenous reference genes and calculated using the comparative Ct method.

Primer sequences used for RT-qPCR are list below.

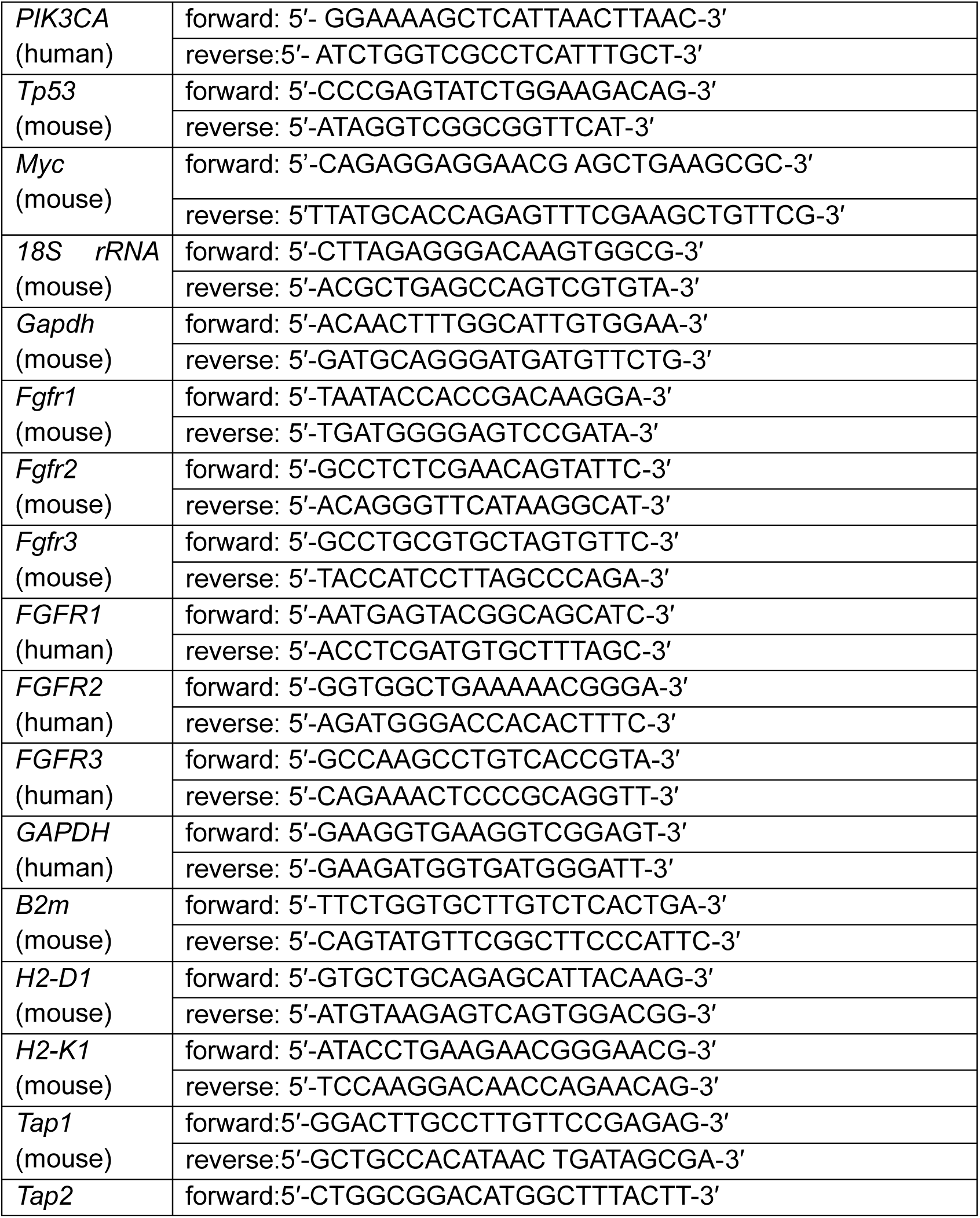
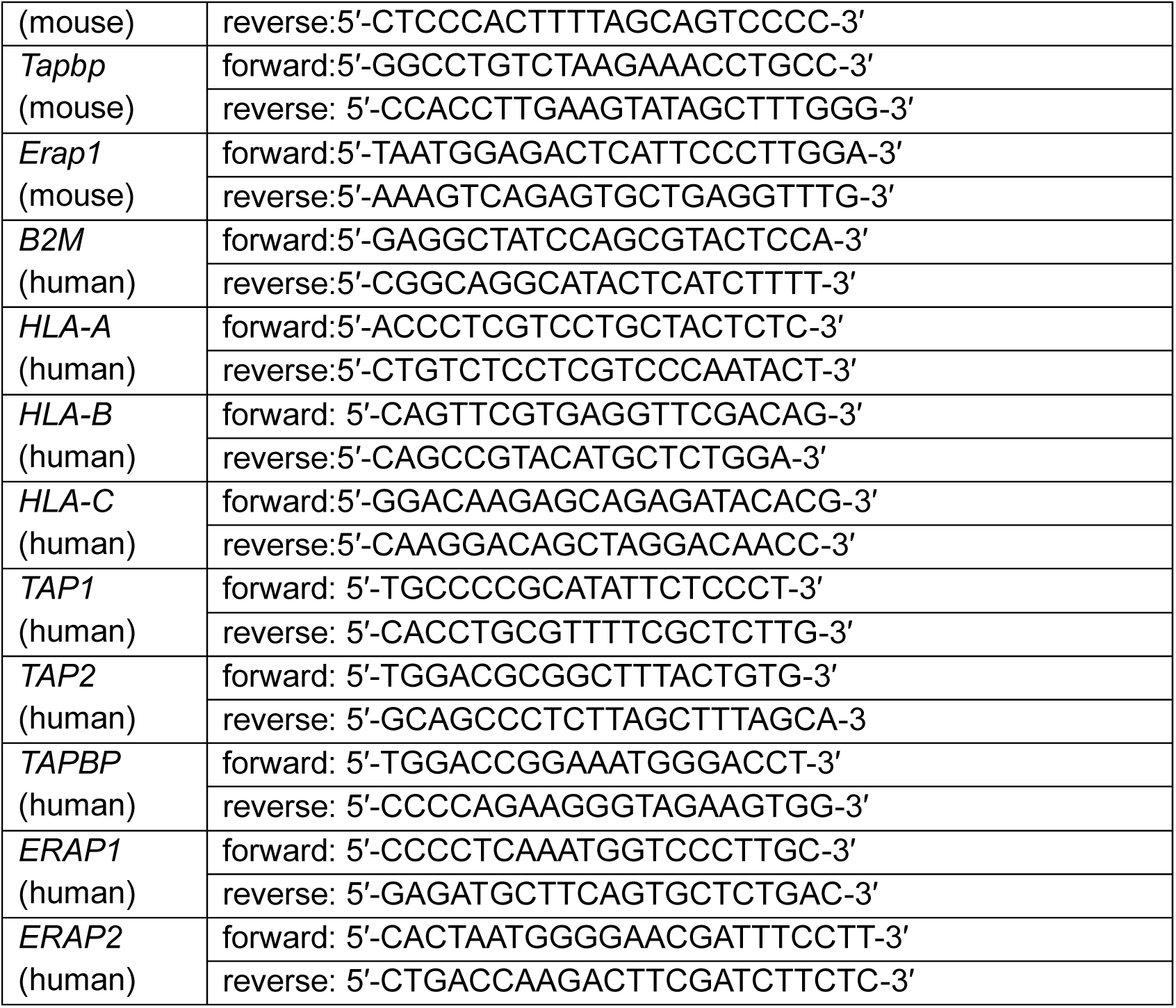

### Western blotting

Western blotting was performed using standard protocols. Primary antibodies used were as follows: HA (3F10, Roche), phospho-S6 ribosomal protein (Ser235/236, Cell Signaling Technology, #2211), phospho-S6 ribosomal protein (Ser240/244, Cell Signaling Technology, #2215), total S6 ribosomal protein (Cell Signaling Technology, #2217), MYC (Santa Cruz Biotechnology, sc-40), phospho-AKT (Ser473, Cell Signaling Technology, #4060), phospho-AKT (Thr308, Cell Signaling Technology, #2965), total AKT (Cell Signaling Technology, #9272), phospho-FRS2 (Tyr436, Cell Signaling Technology, #3861), total FRS2 (R&D Systems, MAB4069), phospho-ERK1/2 (Thr202/Tyr204, Cell Signaling Technology, #9101), total ERK1/2 (Cell Signaling Technology, #9102), phospho-PRAS40 (Cell Signaling Technology, #2997), total PRAS40 (Cell Signaling Technology, #2691), phospho-p70S6 (Cell Signaling Technology, #9205), total p70S6 (Cell Signaling Technology, #9202), FGFR1 (Cell Signaling Technology, #9740), FGFR2 (Cell Signaling Technology, #23328), FGFR3 (Cell Signaling Technology, #4574), phospho-4EBP1(Ser65, Cell Signaling Technology, #9451), phospho-4EBP1 (Thr37/46, Cell Signaling Technology, #2855), total 4EBP1 (Cell Signaling Technology, #9452), phospho-eIF4B (Ser422, Cell Signaling Technology, #3591), total eIF4B (Cell Signaling Technology, #3592), phospho-STAT3 (Tyr705, Cell Signaling Technology, #9145), total STAT3 (Cell Signaling Technology, #9139), and vinculin (Sigma-Aldrich, V9131).

### Flow cytometry

Single-cell suspensions were prepared as described in the “Tissue Digestion” section. Cells were stained with LIVE/DEAD Fixable Aqua Dead Cell Stain (Thermo Fisher Scientific) in cold FACS buffer (PBS supplemented with 0.2% BSA and 5 mM EDTA) for 10 min on ice. Cells were then incubated with anti-mouse CD16/32 (1:50, clone 93, BioLegend) for 15 min on ice to block Fc receptors. Following Fc blocking, cells were stained with fluorophore-conjugated antibodies diluted in FACS buffer for 30 min on ice.

The following antibodies were use: CD45 (1:100, clone 30-F11, BioLegend), CD3 (1:100, clone 145-2C11, BioLegend), CD4 (1:100, clone RM4-5, BioLegend), CD8A (1:100, clone 53-6.7, BioLegend), CD44 (1:100, clone IM7, BioLegend), CD62L (1:100, clone MEL-14, BioLegend), FOXP3 (1:100, clone MF-14, BioLegend), IFN-γ (1:100, clone XMG1.2, BioLegend), TNF-α (1:100, clone MP6-XT22, BioLegend), KLRG1 (1:100, clone 2F1/KLRG1, BioLegend), H2-K^q^ (1:00, clone KH114, BioLegend), HLA-A/B/C (1:100, clone W6/32, BioLegend), β 2-microglobulin (1:100, clone A17082A, BioLegend), CD11b (1:100, clone M1/70, BioLegend), F4/80 (1:100, clone BM8, BioLegend), CD206 (1:100, clone MMR, BioLegend), FGFR1 (1:50, clone M19B2, Thermo Fisher Scientific), FGFR2 (1:50, clone SP273, abcam), and FGFR3 (1:50, clone 2H10B4, Thermo Fisher Scientific).

For detection of unconjugated primary antibodies, cross-adsorbed goat anti-rabbit IgG (H+L) and goat anti-mouse IgG (H+L) secondary antibodies (1:5000; Life Technologies, #A11008 and #A21235) were used.

For intracellular staining, cells were fixed and permeabilized using the Foxp3/Transcription Factor Staining Buffer Set (eBioscience, #00-5523-00) according to the manufacturer’s instructions. For intracellular cytokine analysis, cells were stimulated with Leukocyte Activation Cocktail containing the protein transport inhibitor Brefeldin A (BD Biosciences, # 550583) at 37 °C for 4 hours prior to staining.

Flow cytometry data were acquired on an LSR Fortessa HTS analyzer (BD Biosciences) using BD FACSDiva software and analyzed with FlowJo (BD Biosciences).

### Transcriptome Analysis

Transcriptomic profiling was performed using a custom Ion AmpliSeq panel comprising 4,604 cancer- and immune-associated genes, designed with the Ion AmpliSeq Designer (Thermo Fisher Scientific) as previously described. For each sample, 10 ng of total RNA was used for cDNA library preparation. Libraries were multiplexed, amplified using the Ion OneTouch™ 2 System, and sequenced on an Ion Torrent Proton™ System (Thermo Fisher Scientific). Raw sequencing data were processed using Torrent Suite software and the AmpliSeqRNA analysis plugin (Thermo Fisher Scientific) to generate gene-level count data.

For downstream analyses, including Gene Set Enrichment Analysis (GSEA) and pathway enrichment analysis based on the Kyoto Encyclopedia of Genes and Genomes (KEGG), genes with a mean fold change greater than 2 or less than 0.5 were selected.

### Organotypic tumor spheroid preparation and microfluidic culture

Tumor samples were collected and processed under Institutional Review Board (IRB)-approved protocols at the Dana-Farber/Harvard Cancer Center, with written informed consent obtained from all participants. Patient-derived organotypic tumor spheroids (PDOTS) were generated and cultured as previously described ^65^. For the experiments, PDOTS were exposed to FGFR inhibitors for 7 days to model sustained pathway inhibition under ex vivo conditions prior to downstream analyses.

### Single- nucleus GEM generation and gene expression library construction

Nuclei isolation was performed as previously described ^66^, using low-retention microcentrifuge tubes (Fisher Scientific) to minimize nuclei loss. Briefly, tumor tissues were manually dissociated with fine spring scissors for 10 minutes, homogenized in TST buffer, filtered through a 30-μm MACS SmartStrainer (Miltenyi Biotec), and pelleted by centrifugation at 500 × g for 10 minutes at 4°C. Nuclei pellets were further purified using Anti-Nucleus MicroBeads (Miltenyi Biotec, #130-132-997) following the manufacturer’s instructions.

Purified nuclei were resuspended in 100 μL resuspension buffer (1X PBS supplemented with 1% BSA). Dual-fluorescent staining was performed using ViaStain^TM^ AOPI Staining Solution (Nexcelom Bioscience, #CS2-0106). Nuclei quality and concentration were assessed using a Cellometer-K2 Fluorescent Cell Counter (Nexcelom Bioscience). Although partial loss of nuclei integrity was observed, overall nuclei yield and quality were sufficient for downstream processing.

Single-nucleus gel bead-in-emulsion (GEM) generation was performed using the Chromium X system (10X Genomics) according to the Chromium Next GEM Single Cell 5’ Reagent Kits v2 (Dual Index) protocol. Sequencing-ready gene expression libraries were generated following the standard 10X Genomics 5′ workflow.

### snRNA-seq data analysis

#### A. Dataset quality control

snRNA-seq data were processed using the Cell Ranger pipeline (version 7.1.0, 10x Genomics)^67^. Reads were aligned to the mouse reference genome mm10 (refdata-gex-mm10-2020-A), followed by barcode assignment and UMI counting using the cellranger count workflow. Filtered feature-barcode matrices were imported into Scanpy (version 1.10.0) for downstream analysis ^68^. Doublets were identified and removed using Scrublet (version 0.2.3) ^69^. Nuclei with ≥500 detected genes, mitochondrial gene content <20%, and hemoglobin gene content <5% were retained for further analysis.

#### B. Cell clustering

Normalization, scaling, dimensionality reduction, clustering, and differential gene expression analyses were performed using Scanpy (version 1.10.0). Total-count normalization and log transformation were applied using “normalize_total”, followed by data scaling with “scale”. The top 2,000 highly variable genes were identified using “highly_variable_genes” and used for principal component analysis (PCA). Batch effects were corrected using “Harmony” applied to principal components ^70^. A k-nearest neighbor graph was constructed using the first 30 principal components, and Uniform Manifold Approximation and Projection (UMAP) was used for visualization ^71^. Differentially expressed genes (DEGs) were identified using the “rank_genes_groups” with default parameters.

#### C. Cell annotation

Cell annotation was assigned based on curated marker gene sets refined from published references and validated across multiple datasets ^72, 73^. Epithelial cells were identified by expression of *Cd9*, *Cd24a*, *Sox9*, *Cdh1*, and *Epcam*; fibroblasts by *Pdgfrb*, *Fap*, *Lum*, and *Dcn*; and endothelial cells by *Cdh5* and *Pecam1*. T-cell subsets were classified as CD4⁺ naïve (*Cd4*, *Tcf7*, *Il7r*, *Ccr7*), CD8⁺ naïve (*Cd8a*, *Cd8b1*, *Tcf7*, *Il7r*, *Ccr7*, *Sell*), CD4⁺ exhausted (*Cd4*, *Ctla4*, *Pdcd1*, *Havcr2*, *Lag3*), CD8⁺ exhausted (*Cd8a*, *Cd8b1*, *Ctla4*, *Pdcd1*, *Havcr2*), CD4⁺ regulatory (*Foxp3*, *Il2ra*, *Tnfrsf4*, *Itgae*), and CD8⁺ cytotoxic (*Cd8a*, *Cd8b1*, *Gzma*, *Gzmb*, *Prf1*, *Mki67*). Myeloid populations were annotated as M1 macrophages (*Il1a*, *Il1b*, *Tnf*, *Cxcl10*, *Nos2*), M2 macrophages (*Apoe*, *Cd163*, *Arg1*, *Irf4*), and dendritic cells (*Itgax*, *Cd209a*, *Zbtb46*, *Flt3*). Additional immune subsets included natural killer (NK) cells (*Klrd1*, *Cd94*). Final cell identities were determined by integrating canonical marker expression with unsupervised clustering to ensure consistency across samples.

#### D. RNA velocity analysis

RNA velocity analysis was performed to infer dynamic cell state transitions. Spliced and unspliced transcript counts were generated using velocyto (version 0.17.17) in run10x mode^74^. Velocity moments were computed using scVelo (version 0.3.2) ^75^, and gene-specific splicing kinetics were inferred using *recover_dynamics*. RNA velocities were estimated in *dynamical* mode and visualized using partition-based graph abstraction (PAGA) ^76^.

#### E. Survival analysis

Survival analyses were performed using the R packages survival (version 3.5.5) and survminer (version 0.4.9).

### Statistical analysis

Quantitative data are presented as means ± standard error of the mean (SEM). Statistical analyses were performed using unpaired two-tailed Student’s t-test for comparisons between two groups and one-way analysis of variance (ANOVA) for comparisons among three or more groups, as indicated in the figure legends. All statistical analyses were conducted using GraphPad Prism (version 8.0.2). A p-value< 0.05 was considered statistically significant.

## Data availability

The transcriptomic, snRNA-seq, and genome sequencing data generated in this study have been deposited in the Gene Expression Omnibus (GEO) under accession numbers GSE292814 and GSE311373. Publicly available datasets analyzed in this study are available in GEO under accession numbers GSE225691. Source data are provided with this paper.

## Acknowledgements

We thank Drs. J.A. Sanai, G.B. Mills, and D. Chowdhury for kindly providing EC cell lines. We also thank the Center for Cancer Genomics team for their assistance with snRNA sequencing. This work was supported by grants from the National Cancer Institute (NCI) under award number 1P01CA269021-01A1 (G.I.S., J.L., P.K., U.M., and J.J.Z.), and from the National Institutes of Health (NIH) under award numbers R35CA210057 (J.J.Z.), 1P01CA236749 (G.J.F.) and AI056299 (G.J.F.). K.C. was supported by a start-up fund provided by Boston Children’s Hospital. This work was also supported in part by the Expect Miracles Foundation and the Robert A. and Renée E. Belfer Family Foundation.

## Author Contributions

X.C. and J.J.Z. conceived and designed the study. X.C. performed the majority of the experiments. E.H., G.A.,Y.D.Z., and K.F.C. assisted with snRNA-seq data analysis. C.L.Q., R.L.J., Z.W., and B.K. contributed in vitro experiments, and T.J. supported in vivo animal studies. H.G. and S.Z.X. performed transcriptomic data analysis. J.N. provided the TetO-PIK3CA^H1047R^ mouse model, and E.I. provided human EC patient-derived organoids. M.N. reviewed histological specimens. G.J.F. generously provided the anti-PD-1 antibody. X.C. and J.J.Z. analyzed the data and wrote the manuscript, with G.I.S., P.K., J.L., G.J.F, and U.M. reviewing and editing the manuscript and offering critical feedback.

## Competing interests

G.J.F. has patents/pending royalties related to the PD-L1/PD-1 pathway from Roche, Merck MSD, AstraZeneca, Bristol-Myers-Squibb, Merck KGA, Boehringer-Ingelheim, Dako, Leica, Mayo Clinic, Eli Lilly, and Novartis. G.J.F. has served on advisory boards for iTeos, NextPoint, IgM, GV20, IOME, Bioentre, Santa Ana Bio, Simcere of America, and Geode, and holds equity in Nextpoint, iTeos, IgM, Invaria, GV20, Bioentre, and Geode. G.I.S receives research funding from Merck KGaA/EMD Serono, Artios, Eli Lilly, and Pfizer, and has served on advisory boards for Merck KGaA/EMD Serono, Circle Pharmaceuticals, Concarlo Therapeutics, Schrodinger, FoRx Therapeutics, and MycRx. G.I.S holds patents entitled “Dosage regimen for sapacitabine and seliciclib” and “Compositions and Methods for Predicting Response and Resistance to CDK4/6 inhibition”. J.J.Z. is a co-founder and director of Crimson BioPharma Inc. and Geode Therapeutics Inc. All these activities are not related to this work. The remaining authors declare no competing interests.

## Notes

### Competing Interest Statement

J.J.Z. is a co-founder and director of Crimson BioPharma Inc. and Geode Therapeutics Inc. (not related to this work). G.J.F. has patents and pending royalties related to the PD-L1/PD-1 pathway from Roche, Merck MSD, AstraZeneca, Bristol-Myers Squibb, Merck KGaA, Boehringer Ingelheim, Dako, Leica, Mayo Clinic, Eli Lilly, and Novartis. G.J.F. has served on advisory boards for iTeos, NextPoint, IgM, GV20, IOME, Bioentre, Santa Ana Bio, Simcere of America, and Geode, and holds equity in NextPoint, iTeos, IgM, Invaria, GV20, Bioentre, and Geode (not related to this work). G.I.S. receives research funding from Merck KGaA/EMD Serono, Artios, Eli Lilly, and Pfizer. He has served on advisory boards for Merck KGaA/EMD Serono, Circle Pharmaceuticals, Concarlo Therapeutics, Schrodinger, FoRx Therapeutics, and MycRx. He holds patents entitled "Dosage regimen for sapacitabine and seliciclib" and "Compositions and Methods for Predicting Response and Resistance to CDK4/6 inhibition" (not related to this work).

