## Supplemental Fig1-7 for "Targeting FGFR signaling overcomes therapeutic resistance and immune evasion in oncogenic PIK3CA-driven serous-like endometrial cancer"

### Supplementary Fig. 1

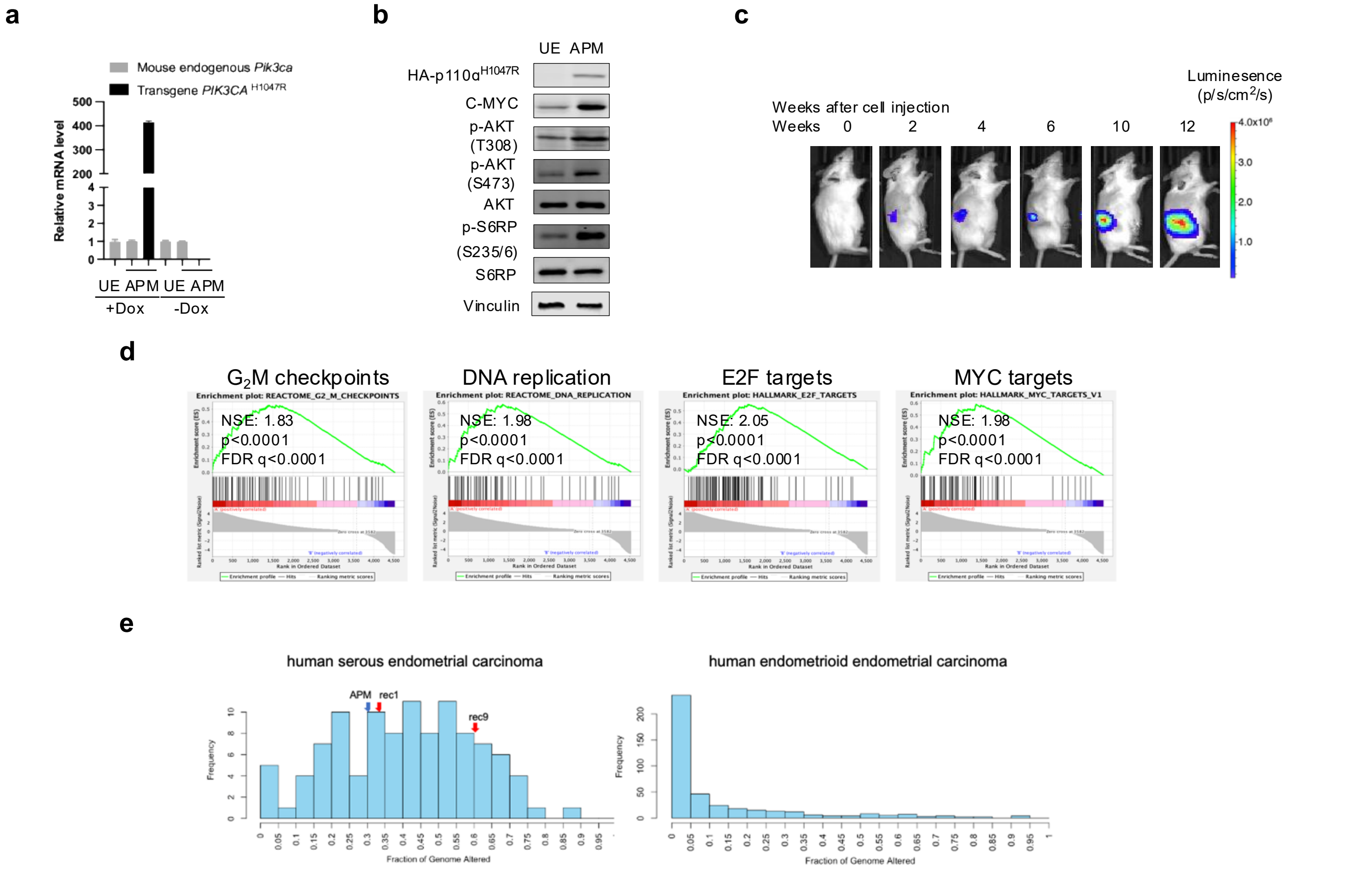


| | | APM | | Rec1 | | Rec9 | |
|---|---|---|---|---|---|---|---|
| Method | Total numbers of bases | Number of CNV | FGA | Number of CNV | FGA | Number of CNV | FGA |
| CNVkit (exome) | 912677426 | 299641482 | 0.33 | 268929392 | 0.3 | 559340263 | 0.61 |

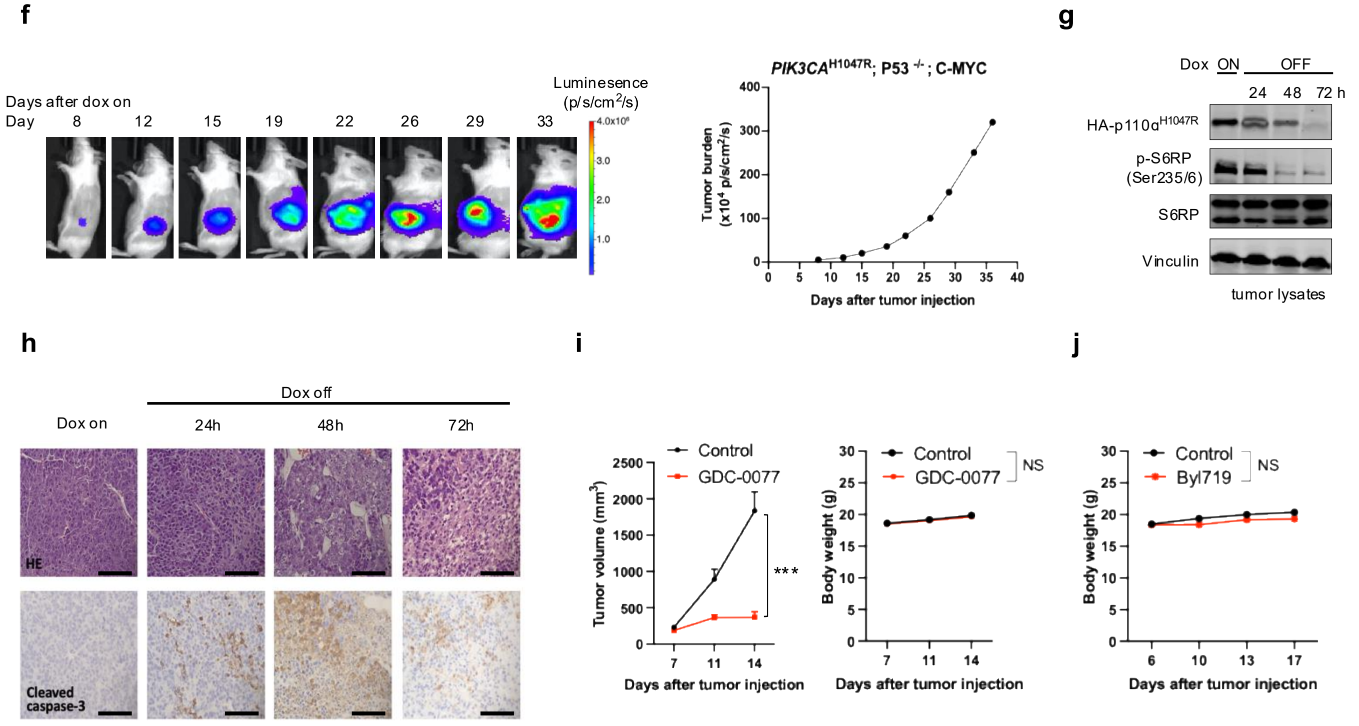

**Supplementary Fig. 1. Establishment and characterization of the PIK3CA$^{H1047R}$; Trp53$^{-/-}$ ; Myc mouse model. (a)** RT-qPCR of the tet-inducible PIK3CA$^{H1047R}$ transgene and endogenous Pik3ca in uterine tissues from tumor-bearing mice with (+) or without (-) doxycycline (n=3 per group). **(b)** Immunoblotting analysis of PIK3CA, c-MYC, and downstream signaling components. UE, uterine surface epithelial (normal) cells. **(c)** Representative in vivo bioluminescence imaging illustrating induction of tet-regulated PIK3CA$^{H1047R}$ expression. **(d)** RNA-seq–based GSEA comparing APM tumors with a published PTEN-loss EC GEMM reveals strong enrichment of G2M checkpoint, DNA replication stress-related programs, E2F targets, and MYC target pathways in APM tumors relative to the PTEN-loss model, consistent with molecular features of human SEC. Data are presented as normalized enrichment scores (NES) with FDR-adjusted q values, based on n = 3 biologically independent tumors per genotype. **(e)** Distribution of FGA in TCGA SEC demonstrates a high burden of CNV (median FGA = 0.41), whereas non-serous TCGA EC samples show a markedly lower alteration burden (median FGA ≈ 0.07). The APM primary tumor model and its recurrent tumors (Rec1 and Rec9) exhibit FGA values of 0.30 and 0.61, respectively, comparable to those observed in human SEC. **(f)** Representative bioluminescence imaging of mice bearing orthotopic, luciferase-expressing APM allografts (n=5). **(g)** Immunoblot analysis of APM tumors with doxycycline on versus after withdrawal. **(h)** H&E staining of APM tumors with doxycycline on versus after withdrawal (scale bar, 50 µm). **(i)** Tumor volume and body weight changes in APM allografts treated with GDC-0077 versus vehicle (n=8-10 per group). **(j)** Body weight trajectories of control- and BYL719-treated mice over time. No treatment-related toxicity or significant body-weight loss was observed.

**Supplementary Fig. 2**

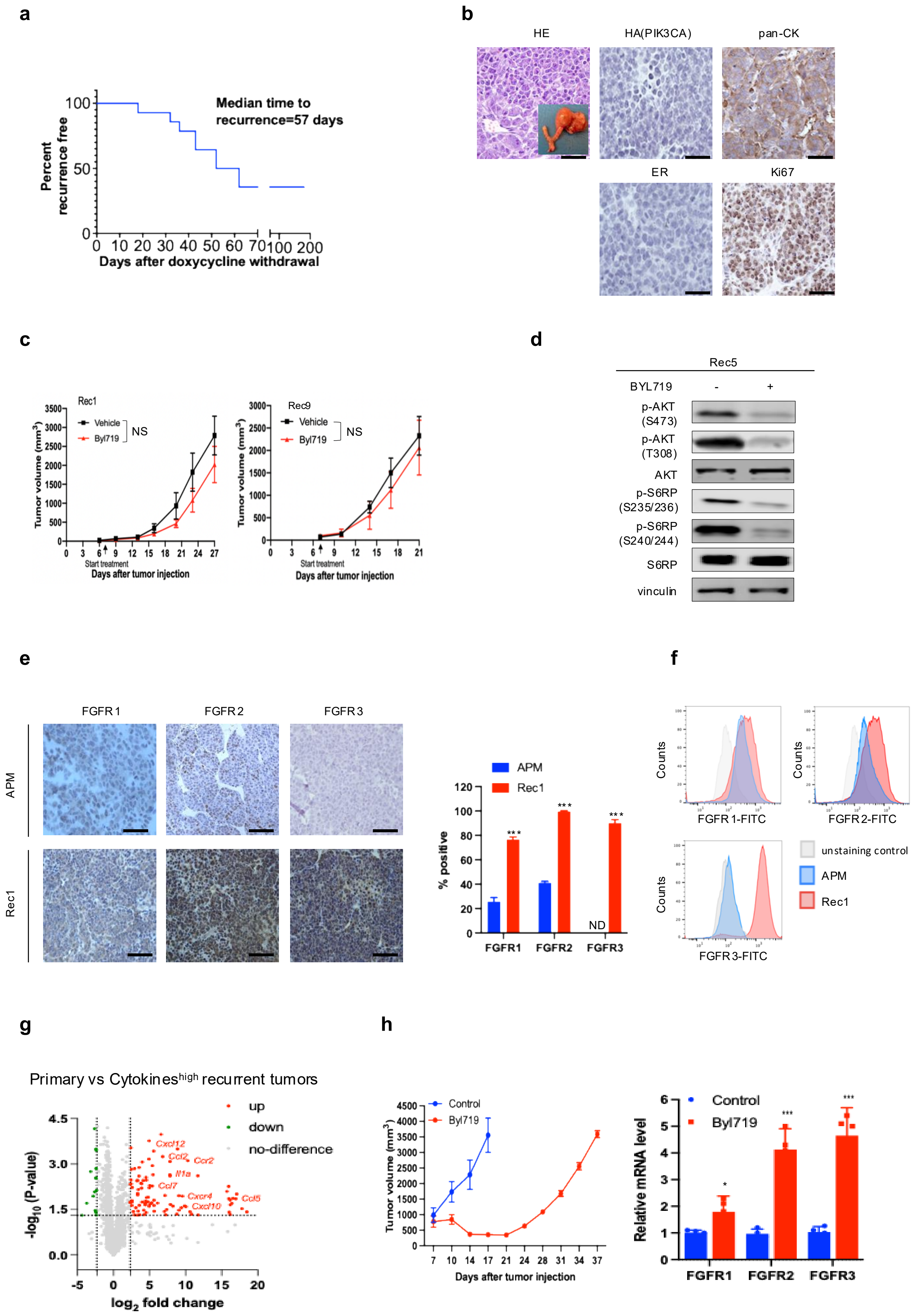

**Supplementary Fig. 2. Characterization of recurrent tumors following PIK3CA$^{H1047R}$ withdrawal. (a)** Time to tumor recurrence following doxycycline withdrawal (n=14). **(b)** Immunohistochemistry analysis of recurrent tumors (scale bar, 50 μm). **(c)** Tumor growth curves of recurrent tumors Rec1 and Rec9 treated with BYL719 (n=8 per group). **(d)** Immunoblot analysis of Rec5 tumor cells treated with BYL719. **(e, f)** Immunohistochemistry (e) and flow cytometric analysis (f) showing FGFR expression in primary versus recurrent tumors. **(g)** Volcano plot illustrating significantly altered gene expression in Cytokines$^{high}$ recurrent tumors based on transcriptomic analysis. (**h**) Tumor growth curves of primary APM tumors treated daily with BYL719 (20 mg/kg; n=8 per group). Right, FGFR expression levels in vehicle-treated control tumors and APM allografts after one month of BYL719 treatment (n=4).

### Supplementary Fig. 3

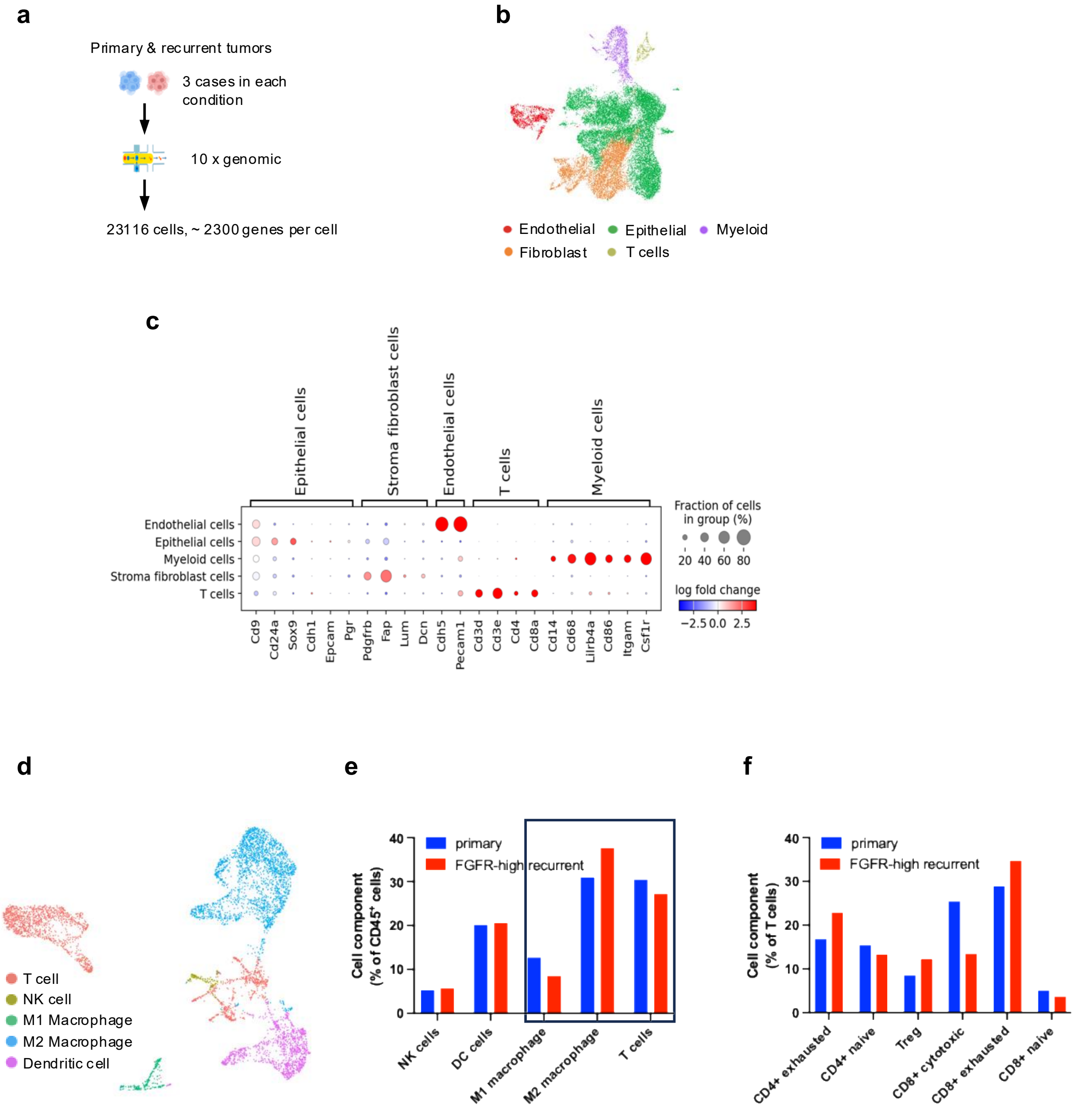


**Supplementary Fig. 3. snRNA-seq analysis of FGFR-high recurrent tumors following *PIK3CA*$^{H1047R}$ withdrawal. (a)** Schematic illustrating the single-nucleus RNA-sequencing workflow for primary and recurrent tumors (n=3 per group). **(b)** UMAP projection showing major cell populations across all samples, including epithelial, fibroblast, endothelial, myeloid, and T cells. **(c)** Expression of canonical marker genes and differentially expressed genes (DEGs) across major cell types. **(d)** UMAP projection of CD45$^{+}$ cells from primary and recurrent tumors, reclustered into T cells, NK cells, M1- and M2-like macrophages, and dendritic cells. **(e)** Quantification of immune cell subsets showing increased M2-like macrophages with concomitant reductions in M1-like macrophages and total T cells in FGFR-high recurrent tumors. **(f)** Composition of T-cell subsets in primary versus recurrent tumors, demonstrating depletion of cytotoxic CD8$^{+}$ T cells and enrichment of exhausted T cells and Tregs in recurrent tumors.

### Supplementary Fig. 4

**a** A panel of human EC lines with known mutations in major oncogenes and tumor suppressors

| | HHUA | EN | SNG-M | MFE-296 | HEC-6 | HEC-1-A | ARK-1 | HEC-151 | HEC-1-B | Nou-1 |
|---|---|---|---|---|---|---|---|---|---|---|
| PIK3CA | R88Q | T1025A | R88Q | P539R | R108H | G1049R | E542K | C420R | G1049R | R38H |
| TP53 | A138V | R273H | R280T | Y220C | R273H | R248Q | R248W | V31I | R248Q | WT |
| PTEN | V290fs | K267fs | V290* | R130Q | V290* | WT | WT | Y76* | WT | no mRNA |
| PIK3R1 | WT | X279-splice | WT | WT | WT | WT | WT | WT | WT | WT |
| KRAS | G12V | WT | G12V | WT | V160A | G12D | WT | WT | D12D | G12D |
| EGFR | WT | WT | WT | WT | A289V | WT | WT | WT | WT | WT |
| HER2-amplified | No | No | No | No | No | No | Yes | No | No | No |
| Pathological subtypes | Endometrial adenocarcinoma | Endometrial carcinoma | Endometrial adenocarcinoma | EEC | Uterine adenosquamous carcinoma | Endometrial carcinoma | SEC | Endometrial carcinoma | Endometrial carcinoma | Endometrial carcinoma |
| BYL719 $IC_{50}$ (μM) | 27 | 16.8 | 12.3 | 12.3 | 5.1 | 3.2 | 2.9 | 2.8 | 2.4 | 0.9 |
| Inavolisib $IC_{50}$ (μM) | 15.2 | 12.9 | 20 | 6.7 | 3.7 | 2.0 | 0.05 | 0.6 | 0.6 | 0.6 |

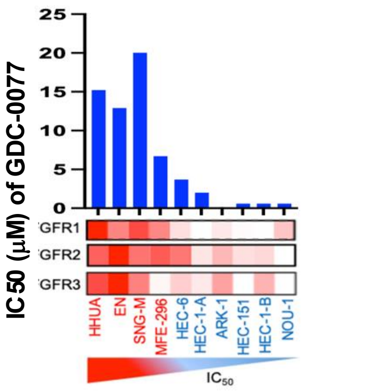


**b**

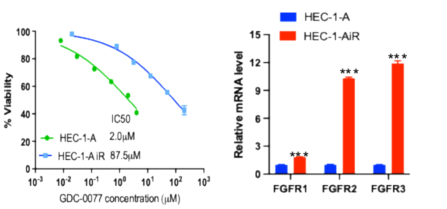


**c**

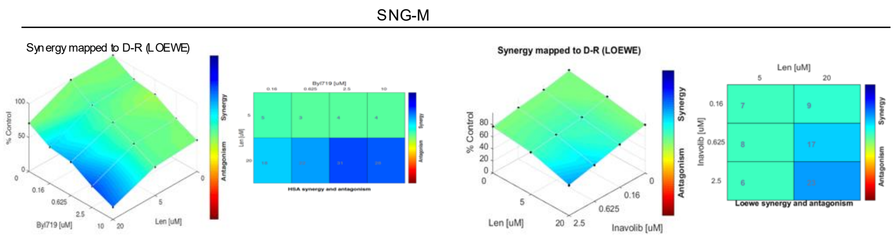


**d**

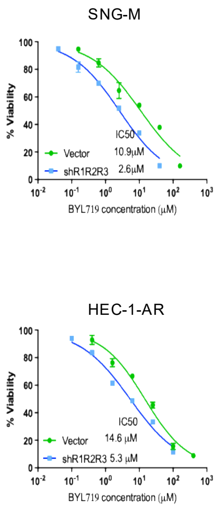


**e**

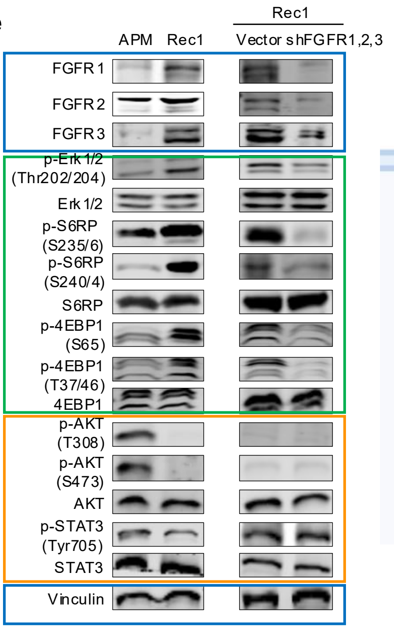


**f**

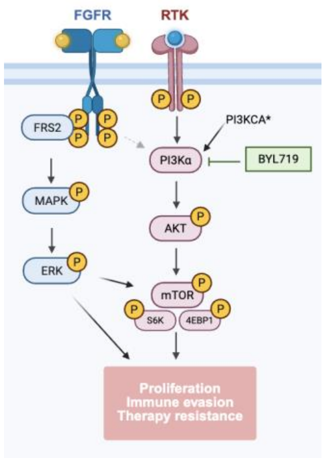


**g**

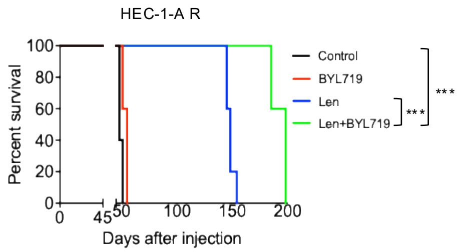


**h**

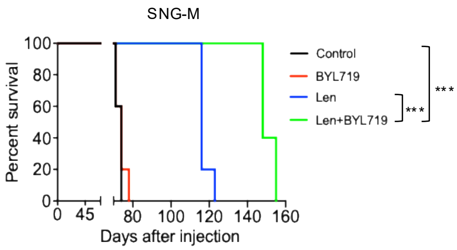


**i**

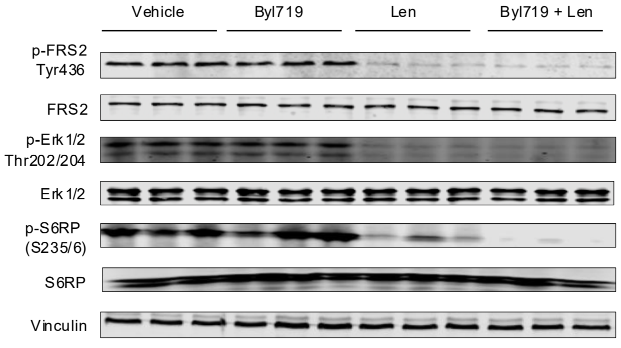

### Supplementary Fig. 4 (continued)

**j**

SNG-M

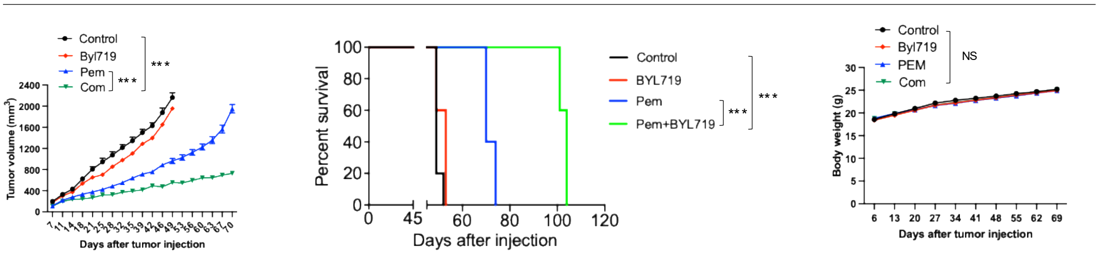


**k**

APM-R

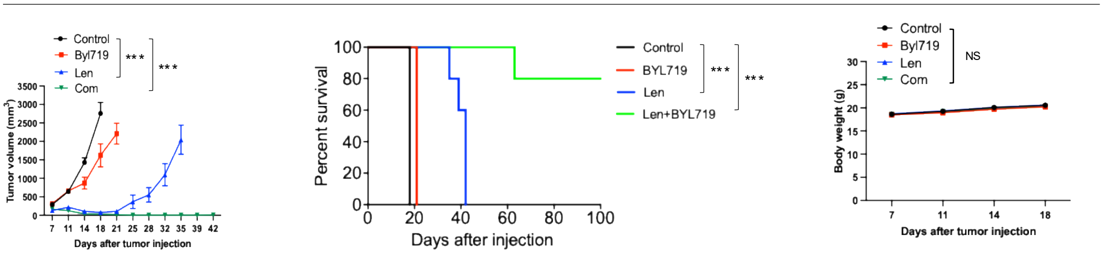


**l**

**HEC-1-A**

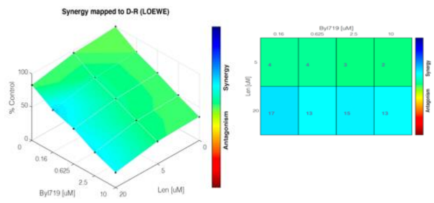


**m**

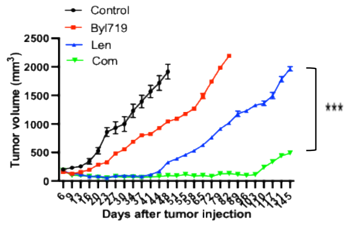

**Supplementary Fig. 4. FGFR signaling limits sensitivity to PI3Kα inhibition. (a)** $IC_{50}$ values for BYL719 and GDC-0077 across ten PIK3CA-mutant human EC cell lines. EEC, endometrioid endometrial cancer. **(b)** Dose-response curves showing sensitivity to inavolisib in HEC-1-A cells (sensitive) and their matched resistant counterpart (left), together with FGFR expression level in the sensitive versus resistant lines (right). **(c)** Isobologram analyses showing additive to synergistic effects of BYL719 plus lenvatinib and inavolisib plus lenvatinib on SNG-M cell viability (synergy-antagonism analysis using Combenefit). **(d)** Effect of FGFR knockdown on BYL719 sensitivity in SNG-M and HEC-1-A R cells. **(e)** Immunoblot analysis of PI3K/AKT, MAPK, and STAT3 signaling in primary tumors, recurrent tumors, and recurrent tumors with FGFR knockdown. **(f)** Schematic summary of the proposed mechanism in which FGFR upregulation sustains mTOR signaling through an AKT-independent, FRS2-MAPK-ERK axis despite PI3Kα inhibition, thereby promoting tumor cell survival, therapeutic resistance, and immune evasion. **(g, h)** Kaplan-Meier survival analyses of HEC-1-A R and SNG-M xenografts treated with lenvatinib and BYL719 (n=5 per group). **(i)** Immunoblot analysis of SNG-M-derived xenografts treated with the indicated therapies. **(j)** Tumor growth curves, Kaplan-Meier survival analyses, and body-weight measurements of SNG-M xenografts treated with pemigatinib plus BYL719 (n=8-10 per group). No treatment-related toxicity or significant body weight loss was observed. **(k)** Tumor growth curves, Kaplan-Meier survival analyses, and body weight measurements of BYL719-resistant APM-R tumors treated with BYL719, lenvatinib, or the combination (n=10 per group). No treatment-related toxicity or significant body weight loss was observed. **(l, m)** Isobologram analysis of BYL719 plus lenvatinib in HEC-1-A cells (diagonal line indicates additivity), and tumor growth curves of HEC-1-A xenografts treated with the indicated therapies (n = 8-10 per group).

### Supplementary Fig. 5

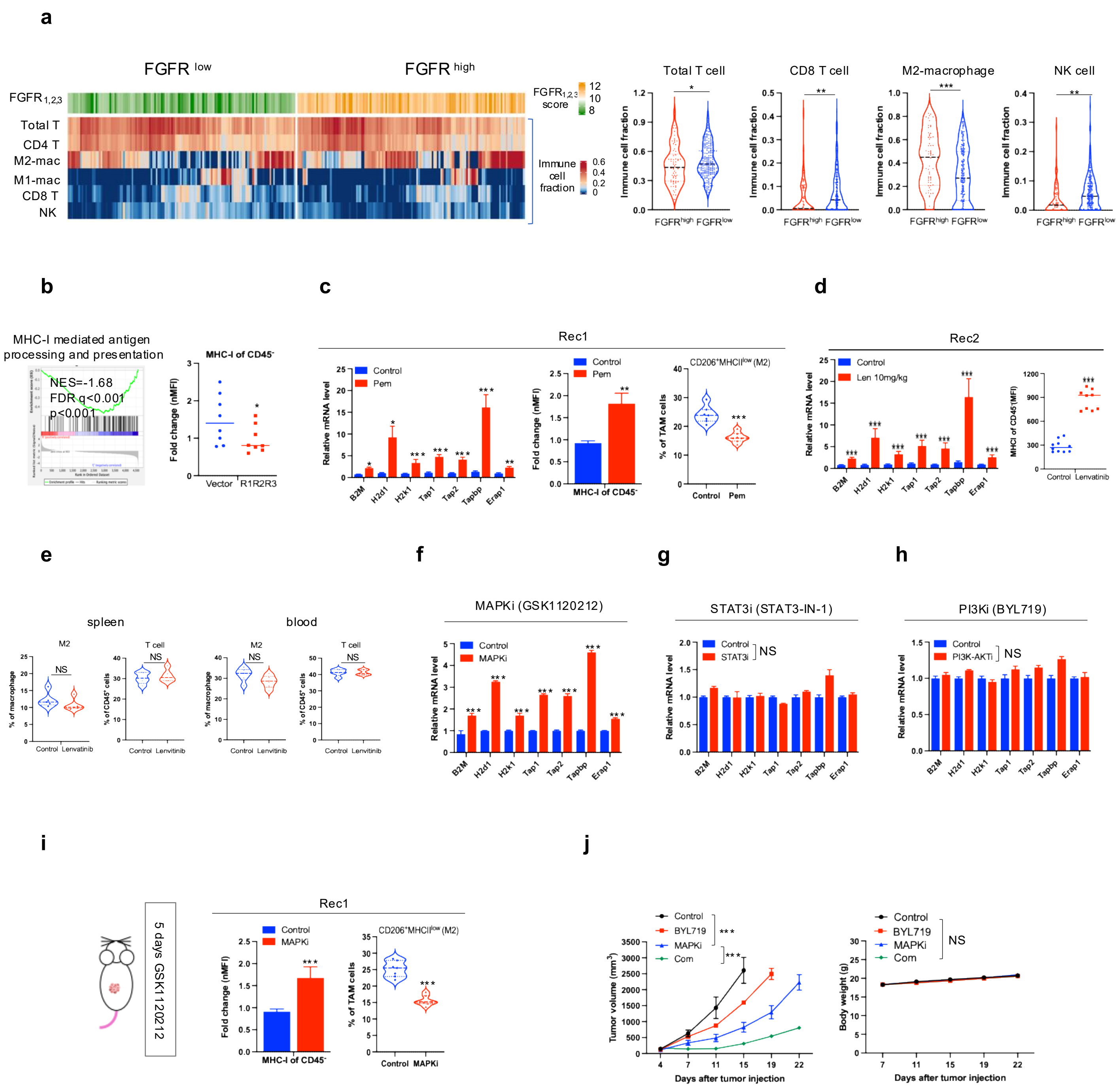

**Supplementary Fig. 5. FGFR inhibition enhances antigen presentation and reduces M2-like TAMs. (a)** Heatmap showing relative FGFR1-3 expression and immune cell composition across uterine cancer samples from the TCGA UCEC cohort. A composite FGFR1-3 score was calculated for each sample as the mean $log_2$ expression of FGFR1, FGFR2, and FGFR3. Immune cell fractions were estimated using CIBERSORT and normalized to sum to 1 per sample. Samples were ordered by increasing FGFR composite score and stratified into FGFR-low and FGFR-high groups defined by the bottom and top quartiles of the FGFR1-3 composite score (n = 66 each; left). Violin plots showing CIBERSORT-estimated immune cell fractions in FGFR-high tumors (top quartile, n = 66) compared with all remaining tumors (FGFR low/others, n=198). FGFR-high tumors exhibit reduced total T cells, $CD8^+$ T cells, and NK cells, together with an increased proportion of M2-like macrophages, consistent with a more immunosuppressive and less inflamed tumor microenvironment (right). **(b)** Downregulation of antigen presentation genes and MHC-I expression in primary tumors overexpressing FGFRs, assessed by GSEA and flow cytometry. **(c, d)** Upregulation of antigen presentation-related genes and reduction of M2-type TAM populations in recurrent tumors following lenvatinib or pemigatinib treatment (n = 8-10 per group). **(e)** Flow cytometric analysis of M2-like macrophages and T cells in the spleen and peripheral blood of naïve mice treated with vehicle or lenvatinib. No significant differences were observed between groups (n = 5 mice per group; NS, not significant), indicating that, in the absence of tumor, lenvatinib has minimal direct impact on systemic immune cell populations. **(f-h)** RT-qPCR analysis of antigen processing and presentation genes in Rec1 tumor cells treated in vitro with inhibitors targeting major FGFR downstream pathways: MAPK/ERK (MAPKi, GSK1120212), STAT3 (STAT3i, STAT3-IN-1), or PI3K-AKT (PI3Ki, BYL719). Only MAPK inhibition robustly upregulates antigen presentation-related genes, whereas STAT3 or PI3K-AKT inhibition has minimal or no effect. **(i)** In vivo validation of MAPK-dependent immune modulation in Rec1 tumors. Flow cytometric analysis of tumors from mice treated with GSK1120212 for 5 days shows increased MHC-I expression (MFI) on $CD45^-$ cells (left) and a reduced percentage of $CD206^+MHCII^{low}$ M2-type TAMs (right), indicating enhanced antigen presentation and macrophage repolarization. **(j)** Tumor growth curves and body weight trajectories of mice bearing FGFR-high, BYL719-resistant APM-R tumors treated with BYL719, GSK1120212, or the combination (n = 8 per group). Combination therapy produces the greatest inhibition of tumor growth without affecting body weight, supporting MAPK-ERK as a key functional effector downstream of FGFR in driving tumor progression and resistance.

### Supplementary Fig. 6

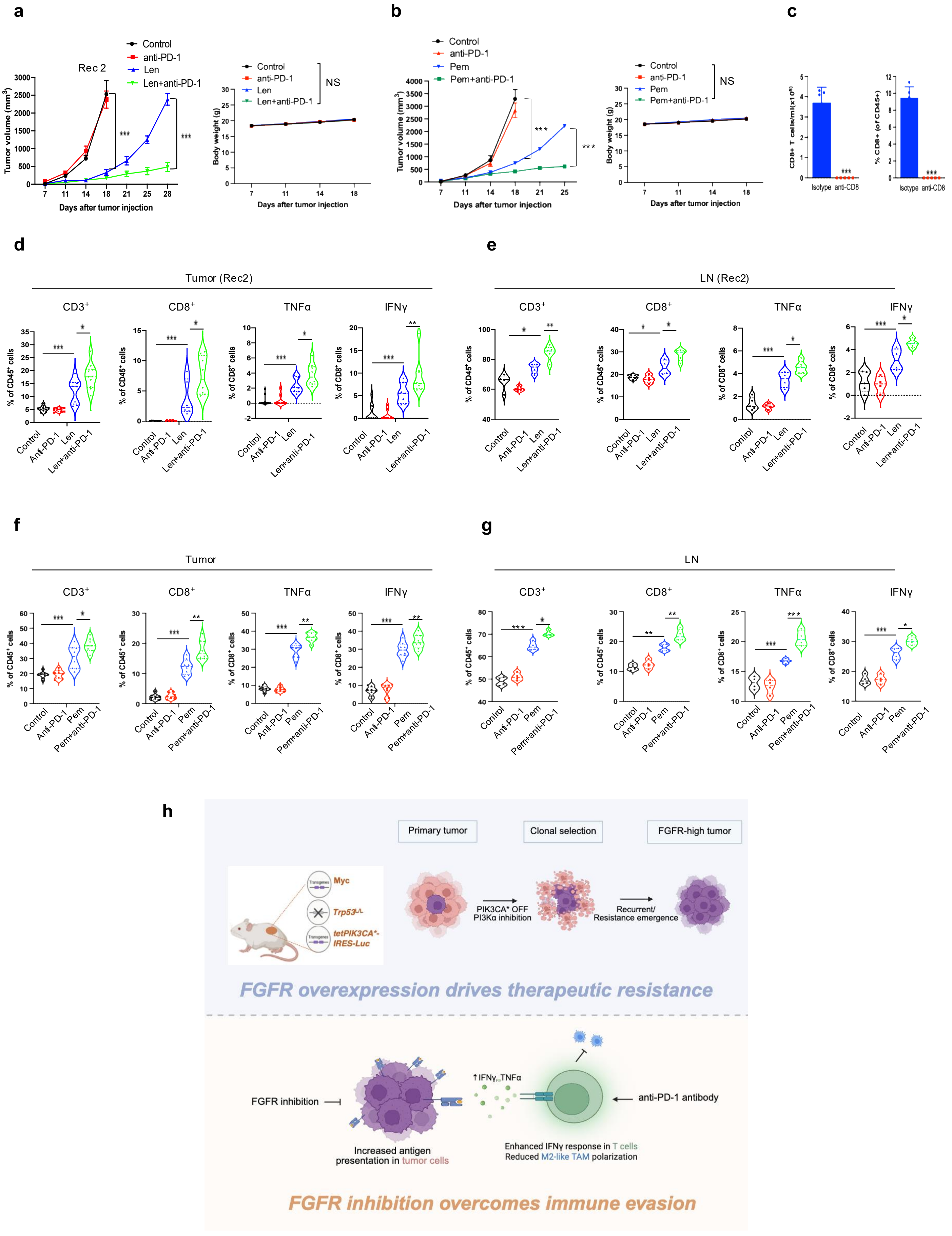

**Supplementary Fig. 6. FGFR inhibition enhances response to immunotherapy and promotes antitumor immunity. (a)** Tumor growth curves and body weight measurements showing the response of recurrent tumor Rec2 to lenvatinib alone or in combination with anti-PD-1 therapy (n=10 per group). No treatment-related toxicity or significant body weight loss was observed. **(b)** Tumor growth curves and body weight measurements showing the response of recurrent tumor Rec1 to pemigatinib alone or in combination with anti-PD-1 therapy (n=10 per group). No treatment-related toxicity or significant body weight loss was observed. **(c)** Flow cytometric analysis of peripheral blood $CD8^+$ T cells following treatment with anti-CD8 or isotype control antibodies. **(d-g)** Flow cytometric analysis of intratumoral and draining lymph node $CD3^+$ and $CD8^+$ T-cell populations and effector cytokine production following the indicated treatments (n=8-10 per group). **(h)** Schematic model summarizing how FGFR signaling promotes therapeutic resistance and immune evasion in PIK3CA-driven serous-like EC by suppressing antigen presentation, limiting $CD8^+$ T-cell infiltration and function, and promoting an immunosuppressive tumor microenvironment.

### Supplementary Fig. 7

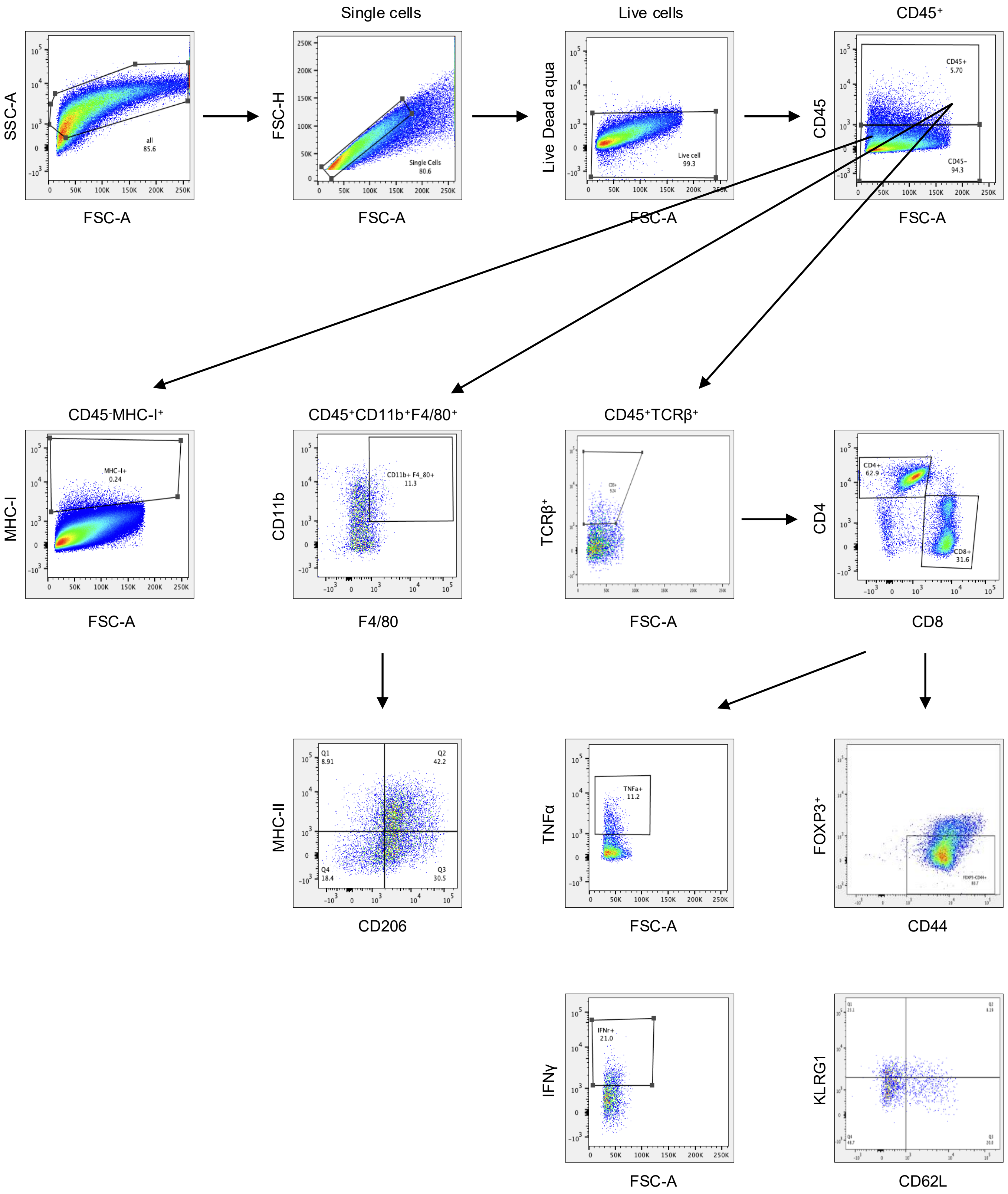


**Supplementary Fig. 7. Gating strategies for flow cytometry.** All flow cytometry plot axes are displayed on a logarithmic scale, except for forward side scatter (FSC), which is shown on a linear scale.
